# Increasing the vitamin C transporter SVCT2 in microglia improves synaptic plasticity and restrains memory impairments in Alzheimer’s disease models

**DOI:** 10.1101/2024.02.29.582679

**Authors:** Camila C. Portugal, Evelyn C. S. Santos, Ana M. Pacheco, Sara Costa-Pinto, Tiago O. Almeida, Joana Tedim-Moreira, Dora Gavin, Teresa Canedo, Fabiana Oliveira, Teresa Summavielle, Sandra H. Vaz, Renato Socodato, João B. Relvas

## Abstract

Alzheimer’s Disease (AD) is characterized by progressive cognitive decline and synaptic dysfunction, often associated with amyloid-beta accumulation and microglial alterations. Here, we investigate the role of the Sodium-dependent Vitamin C Transporter 2 (SVCT2) in microglia to modulate AD-like pathology in mice. Using a combination of RNA sequencing, advanced quantitative proteomics, electrophysiology, behavioral tests, high-throughput imaging, and microglial viral gene delivery, we explore the interplay between SVCT2 expression in microglia, amyloid-beta load, synaptic proteome changes, and synaptic plasticity. Our results demonstrate that SVCT2 expression in microglia decreases with age in the 5xFAD mice, correlating with memory deficits and alterations in synaptic mitochondrial proteome. Importantly, overexpression of SVCT2 in microglia leads to enhanced clearance of amyloid plaques and reconfiguration of the mitochondrial proteome landscape in the synapses, improving synaptic long-term plasticity (LTP) and memory performance. Our findings underscore the SVCT2 overexpression in microglia as a potent strategy to simultaneously decrease amyloid pathology and enhance synaptic plasticity and memory performance, offering new avenues for therapeutic interventions in AD.

## Introduction

Alzheimer’s Disease (AD) is a progressive neurodegenerative disorder and the leading cause of dementia globally. Characterized by progressive memory loss and behavioral changes, AD manifests through diminished decision-making abilities, language difficulties, and personality alterations (Rabinovici, 2019), with anxiety and depression frequently observed as comorbidities (Teri et al., 1999). Histopathologically, AD is marked by reduced brain volume and the presence of amyloid plaques and neurofibrillary tangles. Central to AD’s pathogenesis is oxidative stress, a precursor to other disease hallmarks (Butterfield et al., 2007; Nunomura et al., 2001). It is well documented that plasma antioxidants, including Vitamin C, are depleted in subjects with mild cognitive impairment and AD patients in comparison to healthy subjects (Chang et al., 2014; Rinaldi et al., 2003). Also, there is a significant decrease in ascorbate levels in the cerebrospinal fluid of AD patients in comparison to healthy subjects (Chang et al., 2014; Schippling et al., 2000).

Within this context, the role of Vitamin C is paramount. Ascorbate, the reduced form of vitamin C, is present at high concentrations in the central nervous system (CNS), being crucial for CNS development, functioning, and homeostasis (Portugal, 2024). By exerting its antioxidant functions, ascorbate is oxidized, generating dehydroascorbate, the Vitamin C oxidized form. Due to their chemical structures, their diffusion through the plasma membrane is minimal. Thus, two distinct transport systems mediate the cellular transport of Vitamin C. The reduced form, ascorbate, is taken up by the Sodium Vitamin C co-transporters (SVCT) (Daruwala et al., 1999; Tsukaguchi et al., 1999), while its oxidized form, dehydroascorbate, is transported by the glucose transporters GLUT 1, 2, 3, 4, and 8 (Corpe et al., 2013; Rumsey et al., 2000; Rumsey et al., 1997; Vera et al., 1993).

The high ascorbate concentration in the brain is attributed to the activity of the isoform 2 of the SVCT transporter (SVCT2). This transporter is present in choroid plexus epithelial cells, the route for ascorbate entrance in the CNS (Portugal, 2024). After reaching the brain’s interstitial space, ascorbate is taken up by SVCT2 in neurons, microglia, and oligodendrocytes (Guo et al., 2018; Mun et al., 2006; Portugal et al., 2017). The critical role of the Ascorbate/SVCT2 system in the brain is highlighted by the severe phenotype observed in the SVCT2 KO mice (Sotiriou et al., 2002) and confirmed in a mouse model in which the SVCT2 expression is decreased in the brain (Cao et al., 2023). In both models, the brain shows drastically reduced levels of ascorbate alongside severe and specific brain hemorrhages. Both models die shortly after birth (Cao et al., 2023; Sotiriou et al., 2002). These works highlight the critical role of SVCT2 in maintaining brain ascorbate levels.

Despite the recognized importance of vitamin C in fighting oxidative stress in AD, epidemiological studies have not consistently linked vitamin C supplementation with AD prevention or improvement (Bowman, 2012; Monacelli et al., 2017). This inconsistency may stem from variations in vitamin C bioavailability, influenced by the transport mechanisms governed by SVCT2. Recent data from single-nucleus sequencing of human AD patients reveal decreased SVCT2 expression in microglia associated with AD (Prater et al., 2023), suggesting a compromised capacity for vitamin C uptake in AD microglia. This finding critically shows the necessity of re-evaluating the role of vitamin C in AD, focusing on the transport mechanisms that govern its bioavailability in the brain.

Genetic analyses in diagnosed AD patients highlight the significance of microglial genes as genetic risk factors for the disease (Efthymiou & Goate, 2017), a finding corroborated by animal studies identifying genes modulating microglial responses to amyloid-beta (Aß) plaques (Sala Frigerio et al., 2019). This burgeoning evidence positions microglial homeostasis at the forefront of AD pathology research, underscoring the complex interplay between genetic predispositions and environmental triggers in disease progression. Notably, exposure of microglia to Aß induces a distinct phenotype known as Damage-Associated Microglia (DAM), which is implicated in neuronal damage through the upregulation of inflammatory mediators and reactive oxygen species (ROS) (Keren-Shaul et al., 2017).

Our previous work reveals a pivotal role of SVCT2 in maintaining microglial homeostasis (Portugal et al., 2017). Adult heterozygous SVCT2 mice exhibit significant microgliosis, indicating the critical nature of SVCT2-mediated ascorbate uptake for microglial function in the adult brain (Portugal et al., 2017). Furthermore, impairments in ascorbate uptake resulting from reduced SVCT2 expression are necessary and sufficient to trigger microglial activation. This activation is characterized by NF-κB translocation to the nucleus, elevated production of pro-inflammatory cytokines (TNF, IL-1ß, IL-6), increased expression of iNOS, and ROS generation (Portugal et al., 2017). Moreover, the overexpression of SVCT2 counteracts LPS-induced microglial activation, highlighting the potential therapeutic value of modulating SVCT2 expression and function in AD.

Building on this groundwork, we delve into how SVCT2 impacts microglial function in a mouse model of AD-like pathology and its implications for disease progression. By investigating the interplay between SVCT2 function, vitamin C transport, and microglial dynamics, we aim to shed light on AD pathophysiology and potential therapeutic targets centered on the regulation of microglial SVCT2.

## Results

### Decrease of Microglial SVCT2 Expression Correlates with Cognitive Decline in the 5xFAD Mice

AD is associated with cognitive decline, linked to molecular changes in brain microglia. Central to these microglial changes is the uptake of Vitamin C, mediated by SVCT2 (Portugal et al., 2017). Indeed, decreased transcripts for SVCT2 are found in human microglia associated with AD (Prater et al., 2023). Therefore, these regulatory mechanisms are hypothesized to become compromised in the aging process, particularly within AD. By leveraging the 5xFAD transgenic mouse model of AD (Oakley et al., 2006), we explored the correlation between microglial SVCT2 expression and memory performance.

Using flow cytometry, we evaluated SVCT2 expression in microglia of 5xFAD and WT mice at different time points (Fig. 1A). This analysis provided snapshots of SVCT2 expression at 2, 4, and 6 months of age, revealing a distinct age-dependent decline in SVCT2 fluorescence intensity among the 5xFAD microglia compared to WT, indicative of decreased transporter expression with pathology progression.

**Figure 1:**
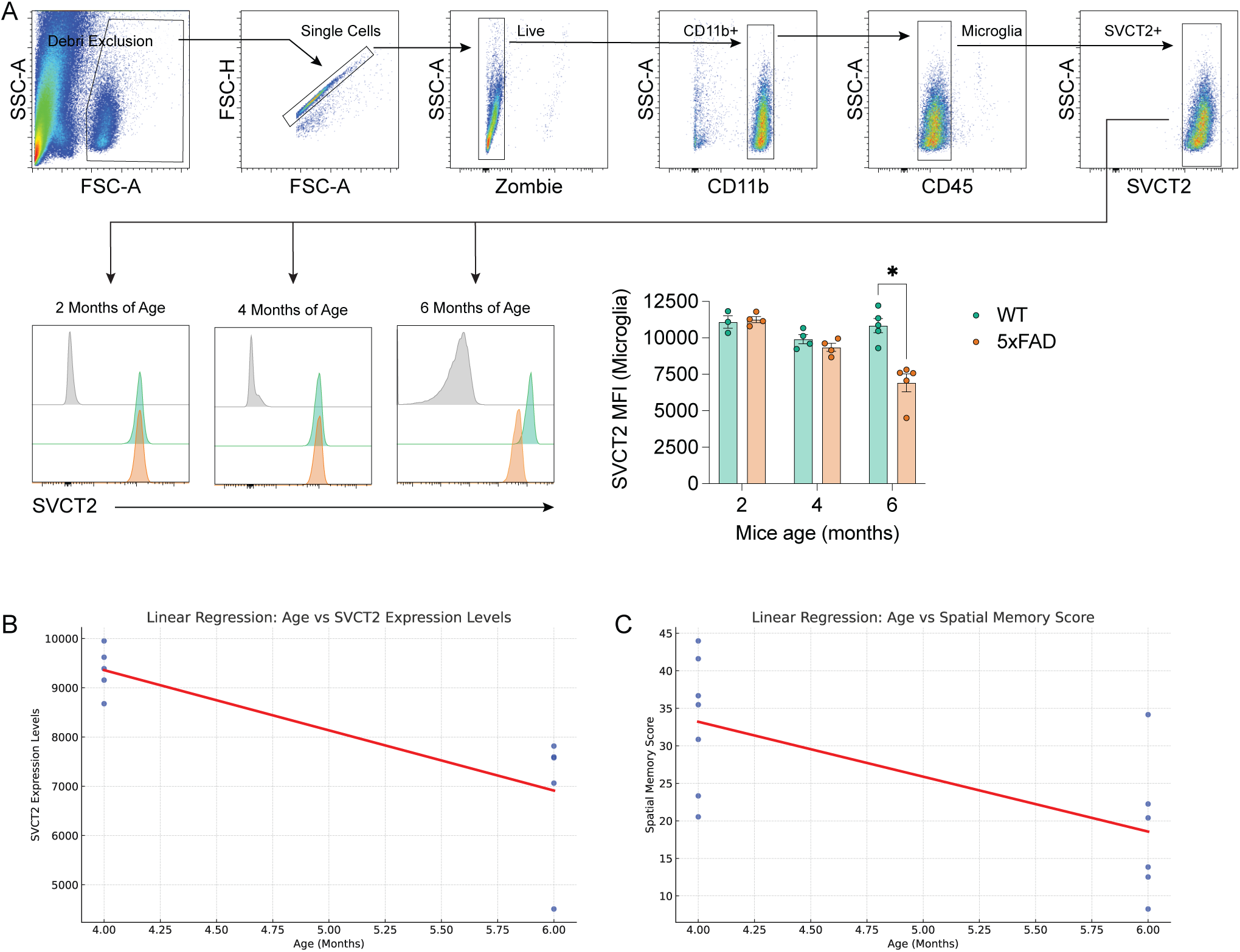
Age-Dependent Decline in SVCT2 Expression in Microglia and Its Correlation with Spatial Memory Deficits in the 5xFAD Mice. A: Flow cytometry analysis of SVCT2 expression in microglia from WT and 5xFAD mice. The gating strategy is depicted, starting with debris exclusion based on forward scatter (FSC-A) and side scatter (SSC-A), followed by identification of single cells, live cells (Zombie negative), CD11b^+^CD45^Dim^ microglia, and then SVCT2^+^ microglia. The histograms represent microglial SVCT2 expression levels at 2, 4, and 6 months of age in both WT (green) and 5xFAD (orange) mice, with the bar graph summarizing the median fluorescence intensity (MFI) of SVCT2 levels in microglia across the different ages (represented by mean ± SEM). *p<0.05 (Two-way ANOVA). Each point in the histograms represents a single mouse. WT mice with 2 months of age (N=3); WT mice with 4 months of age (N=4); WT mice with 6 months of age (N=5); 5xFAD mice with 2 months of age (N=4); 5xFAD mice with 4 months of age (N=5); 5xFAD mice with 6 months of age (N=5) B and C: Panel B depicts the correlation between age and SVCT2 expression levels in microglia, where each point signifies the SVCT2 expression in an individual mouse. Panel C depicts the relationship between age and spatial memory scores, in which each point represents the spatial memory performance of a single mouse.

Next, we utilized regression analyses to further investigate the dynamics of SVCT2 expression over time in the 5xFAD mice, in addition to the snapshot provided by flow cytometry. This statistical approach enabled us to quantitatively model the relationship between age and SVCT2 expression in the 5xFAD mice (Fig. 1B). The significant negative correlation (r = -0.89, p < 0.05) and the slope of the regression line (-1234.6 MFI/month) not only confirmed a decline in microglial SVCT2 but also quantified the rate of this decline over a critical period of 4 to 6 months of age. Alongside this molecular assessment, the spatial memory performance of 5xFAD mice (measured using the Morris water maze test) displayed a similar trend when compared with non-diseased WT littermates within the same timeframe. Regression analysis of the spatial memory data (Fig. 1C) demonstrated a significant decrease in performance scores with age (r = -0.85, p < 0.05), indicating a memory decline over the same period that was consistent with the reduction in microglial SVCT2 expression. These data suggest that the decrease in microglial SVCT2 expression might underpin the memory deterioration associated with AD-like pathology progression in the 5xFAD mice.

### Overexpressing SVCT2 in Microglia Prevents Memory Deficits and Synaptic Plasticity Impairments in the 5xFAD Mice

Building upon the link between decreased microglial SVCT2 expression and memory deficits, we sought to determine the effects of SVCT2 overexpression in microglia on memory performance and synaptic plasticity in the 5xFAD mouse. As a critical part of this work requires the assessment of memory function, all interventions and analyses were performed in the hippocampus. We employed an adeno-associated virus (AAV9) vector containing the minimal CD68 promoter, known for its myeloid cell specificity, to target SVCT2 expression to the microglia within the hippocampus (Rosario et al., 2016). The AAV9 serotype associated with the CD68 promoter successfully induces transgene delivery to microglia, including in the hippocampus (Aschauer et al., 2013; Grace et al., 2016; Grace et al., 2018; O’Carroll et al., 2021). AAV9 vectors harboring either the CD68:mCherry (Control) or CD68:SVCT2-IRES-mCherry (SVCT2) transgenes were administered into the hippocampi of 2-month-old 5xFAD mice *via* stereotactic injections (Fig. 2A), a time point preceding a significant amyloid deposition (Forner et al., 2021). By six months (four months later), a time point selected for its relevance to the emergence of robust AD-like pathology (Oakley et al., 2006), we evaluated the efficiency and specificity of microglial transduction and SVCT2 overexpression (Fig. 2A).

**Figure 2:**
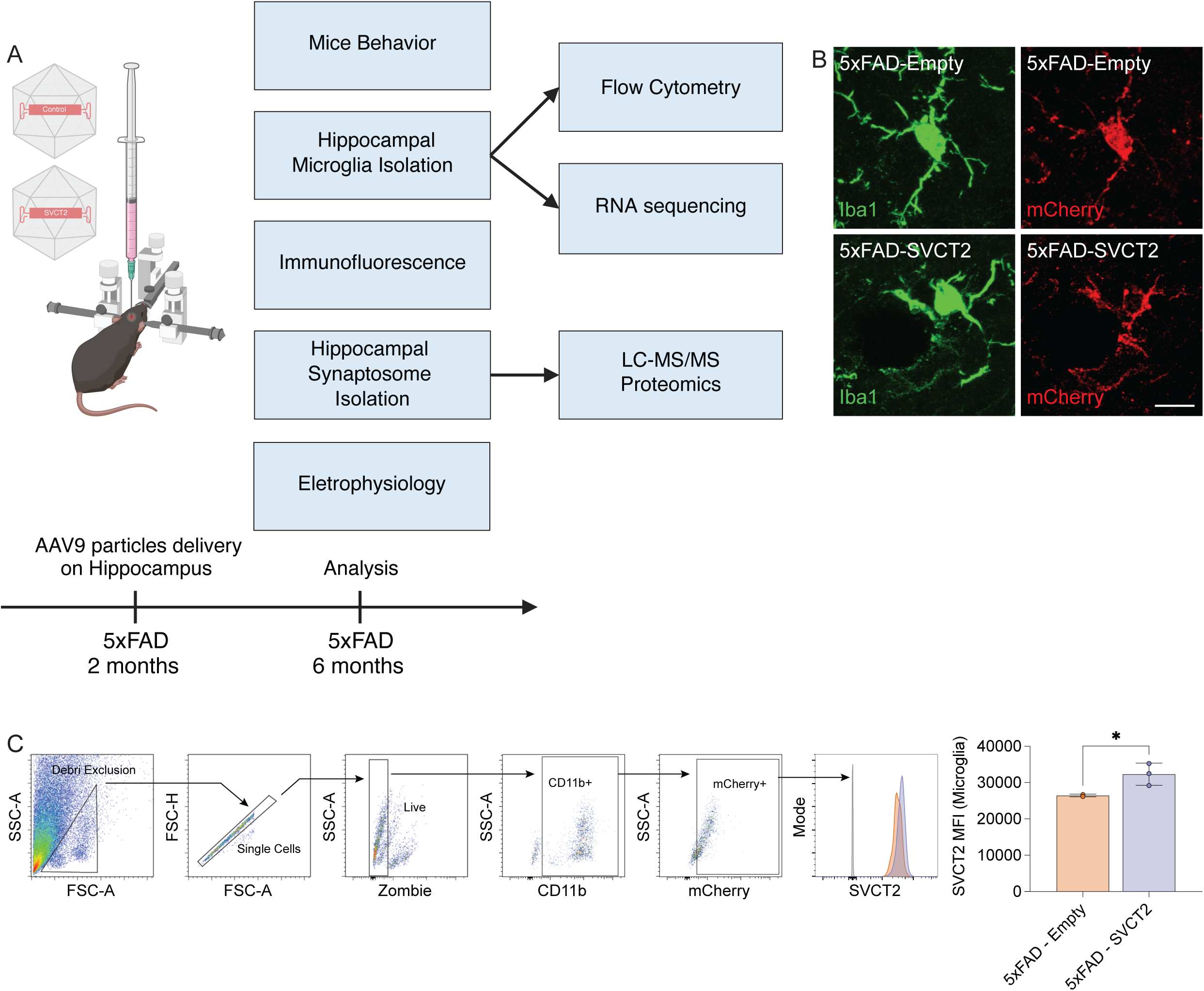
SVCT2 overexpression in hippocampal 5xFAD microglia. A: Schematic representation of the study design showing the stereotactic surgery for the hippocampal delivery of AAV9 particles carrying a minimal CD68 promoter driving mCherry as a reporter. Mice were injected with either AAV9 CD68:mCherry (Empty) or AAV9 CD68:SVCT2:mCherry (SVCT2) at two months of age. Analyses, including behavioral assessment, microglia isolation, immunofluorescence, synaptosome isolation, and electrophysiology, were performed when mice reached six months of age. Subsequent evaluations employed flow cytometry, RNA sequencing, and LC-MS/MS proteomics to investigate the effects of SVCT2 expression on microglial activation and synaptic protein composition. B: Confocal microscopy images displaying hippocampal microglial cells in 5xFAD mice transduced with either Empty or SVCT2 constructs. Microglia are visualized with Iba1 staining (green), and viral delivery is tracked through mCherry immunofluorescence (red). Scale bar = 10 µm. C: Flow cytometry gating strategy analyzing SVCT2 expression in mCherry^+^ microglia. Initial panels show debris exclusion, identification of single cells, and live cell discrimination, followed by the selection of CD11b^+^ microglial cells. The mCherry^+^ population indicates cells transduced with AAV9. SVCT2 expression was assessed within this positively marked population. The bar graph represents the mean fluorescence intensity (MFI), displayed by mean ± SEM, of SVCT2 in mCherry^+^ microglia from both 5xFAD-Empty (N=2) and 5xFAD-SVCT2 (N=3) mice. *p<0.05 (Unpaired t Test)

Confocal-based immunofluorescence analysis revealed notable mCherry fluorescence in hippocampal Iba1^+^ cells, confirming the effectiveness of the CD68 promoter in targeting transgene delivery to microglia (Fig. 2B). Quantitative assessment by flow cytometry identified a substantial mCherry^+^ cell subset, further indicating a robust AAV9-mediated transgene delivery to microglia in 5xFAD animals (Fig. 2C). As expected, 5xFAD mice injected with the SVCT2 virus displayed a significant increase in the SVCT2 expression compared to the mCherry^+^ microglia of 5xFAD animals injected with control AAVs (Fig. 2C), validating the overexpression of SVCT2 in hippocampal 5xFAD microglia.

Following the validation of SVCT2 overexpression in 5xFAD microglia (Fig. 2), we next explored the functional outcomes of this targeted microglial approach on memory performance and synaptic plasticity in the 5xFAD mice. Memory functions were evaluated with the Novel Object Recognition (NOR) test to assess recognition memory and the Morris Water Maze (MWM) to evaluate spatial memory (Fig. 3A). In the NOR test, WT mice injected with the control vector (WT - Empty) showed preference for the novel object, indicating intact recognition memory. In Contrast, 5xFAD mice injected with the control vector (5xFAD - Empty) exhibited a marked reduction in novel object interaction, indicating a recognition memory impairment. Notably, 5xFAD mice overexpressing SVCT2 (5xFAD - SVCT2) demonstrated a significant recovery in novel object preference, approaching levels observed in WT mice (Fig. 3B). Spatial memory was then assessed using the MWM, a well-established assay for hippocampal-dependent memory (Fig. 3C). Path tracking analysis revealed that WT and 5xFAD mice overexpressing SVCT2 showed efficient search patterns and spent a more significant percentage of time in the target quadrant, indicative of successful learning and memory retention. In contrast, 5xFAD mice injected with the control vector exhibited less targeted search patterns and reduced time in the target quadrant, highlighting spatial memory impairment (Fig. 3C).

**Figure 3:**
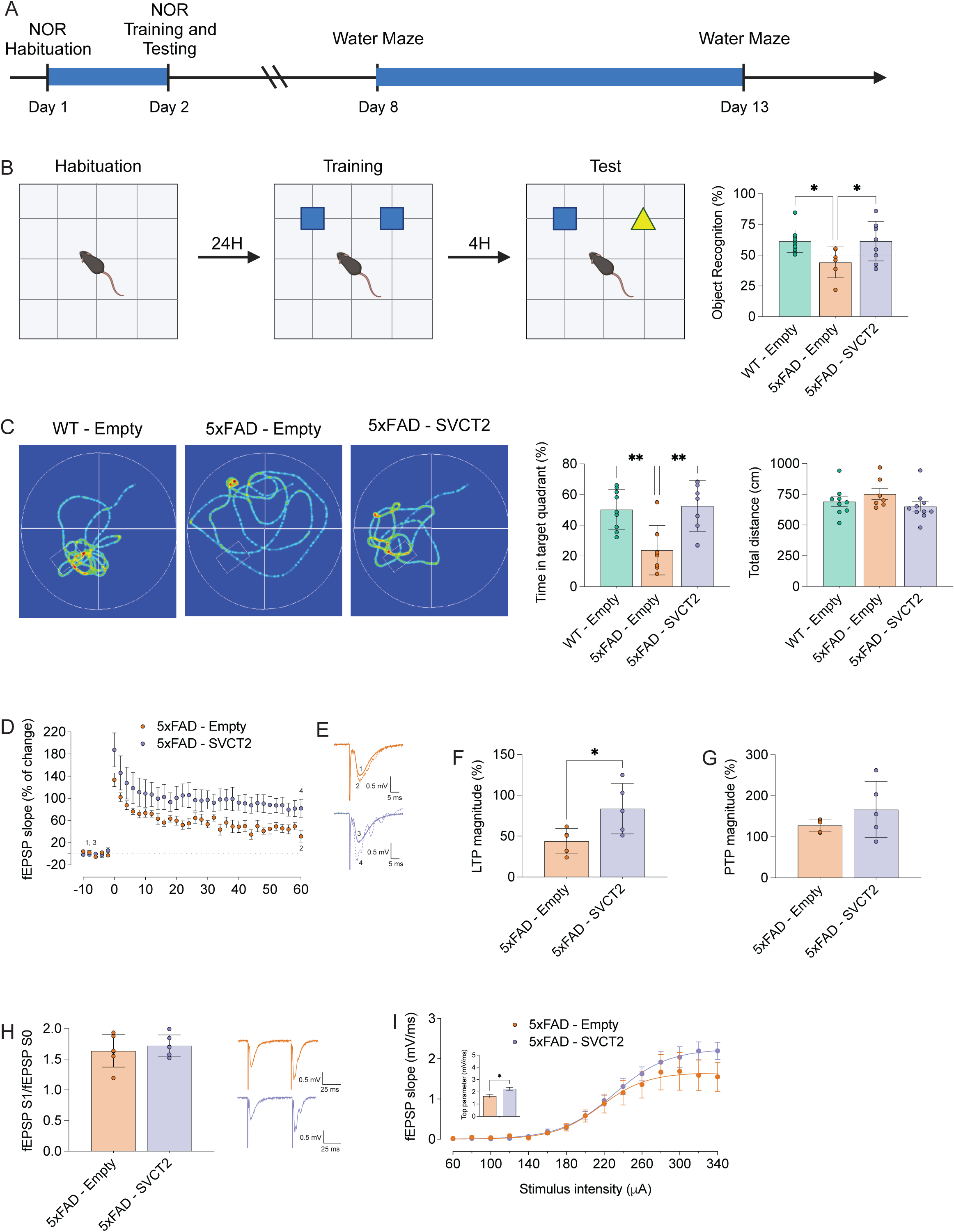
Microglial SVCT2 overexpression improves synaptic function and enhances memory performance in the 5xFAD mice. A: Timeline of the Novel Object Recognition (NOR) and Morris Water Maze (MWM) tests. Mice underwent a habituation phase on Day 1, NOR training and testing on Day 2, followed by MWM training starting on Day 8 and testing on Day 13. B: Schematic representation of the NOR test protocol. The bar graph represents the (%) Recognition, displayed by mean ± SEM, calculated exactly as described in the experimental procedure section. Each point in the histograms represents a single mouse. *p<0.05 (One-way ANOVA) WT-Empty (N=11); 5xFAD-Empty (N=6); 5xFAD-SVCT2 (N=9). C: Left panel: Representative swim paths of wild-type (WT - Empty), 5xFAD mice with the control vector (5xFAD - Empty), and 5xFAD mice overexpressing SVCT2 (5xFAD - SVCT2). Right: Bar graphs representing the (%) time spent in the target quadrant and total distance swam (cm). Both parameters were displayed as mean ± SEM. Each point in the histograms represents a single mouse. **p<0.01 (One-way ANOVA) WT-Empty (N=9); 5xFAD-Empty (N=7); 5xFAD-SVCT2 (N=10). D: SVCT2 overexpressing microglia prevents LTP and basal synaptic transmission deficits in 5xFAD mice. Time course of changes in fEPSP slope after θ-burst stimulation in hippocampal slices from 5xFAD mice with (orange dots) or without (purple dots) microglial overexpression of SVCT2. The graph represents the fEPSP slope (% of change), displayed by mean ± SEM, calculated exactly as described in the experimental procedure section. 5xFAD-Empty (N=5); 5xFAD-SVCT2 (N=5). E: Representative traces of fEPSPs before (bold line) and after (dash line) θ-burst stimulation from hippocampal slices of 5xFAD mice with (orange lines) or without (purple lines) microglial overexpression of SVCT2. Scale bar: 5ms (horizontal), 0.5mV (vertical). F: Comparison of LTP magnitudes obtained in the experiments illustrated in (D). The graph displays mean ± SEM, calculated exactly as described in the experimental procedure section. 5xFAD-Empty (N=5); 5xFAD-SVCT2 (N=5). *p<0.05 (Unpaired t Test) G: Comparison of PTP magnitudes obtained in the experiments illustrated in (D). The graph displays mean ± SEM, calculated exactly as described in the experimental procedure section. 5xFAD-Empty (N=5); 5xFAD-SVCT2 (N=5). H: Data from paired-pulse facilitation (PPF) experiments. Left panel: The bar graph represents the ratio of fEPSP slopes induced by the 2nd (S1) over the first (S0) stimulation (see methods for further details) displayed by mean ± SEM. 5xFAD-Empty (N=5); 5xFAD-SVCT2 (N=5). Right panel: Representative tracings of paired fEPSPs recorded from hippocampal slices of 5xFAD mice with (orange line) or without (purple line) microglial overexpression of SVCT2. Scale bar: 25ms (horizontal), 0.5mV (vertical). I: Input/output (I/O) curves were the fEPSP slope values (ordinates) recorded from hippocampal slices of 5xFAD mice with (purple) or without (orange dots) microglia overexpressing SVCT2 plotted against the intensity of stimulation (abscissae, 60–320 μA). Data are displayed by mean ± SEM. 5xFAD-Empty (N=5); 5xFAD-SVCT2 (N=5). inset: The top parameter of the fitted curves was obtained by the best fit (GraphPad program) of the I/O curves. *p<0.05 (Unpaired t Test)

Synaptic dysfunction is a hallmark of cognitive decline in AD. Hence, we analyzed the impact of microglial SVCT2 overexpression on synaptic plasticity. For this, we evaluated long-term potentiation (LTP) at the CA1 hippocampal synapses in 5xFAD mice overexpressing SVCT2 in microglia or 5xFAD mice injected with control virus. Consistent with our hypothesis, θ-burst stimulation of Schaffer collateral pathways led to an initial surge in the field excitatory postsynaptic potential (fEPSP) slope, which subsequently decreased to a plateau markedly above pre-stimulation baseline values (Fig. 3D). Notably, the LTP magnitude - expressed as the percentage change in fEPSP slope pre- and post-50-60 minutes of θ-burst stimulation - was significantly enhanced in the SVCT2-injected group compared to 5xFAD mice injected with control virus, signifying pronounced facilitation of synaptic strength (Fig. 3D-F). This effect was specific to LTP, as post-tetanic potentiation (PTP) magnitudes remained unaltered between groups (Fig. 3G), underscoring the specificity of SVCT2’s role in modulating long-term forms of synaptic plasticity.

Paired-pulse facilitation (PPF) assay, designed to probe presynaptic alterations, indicated no significant differences in glutamate release probabilities or quantal content between groups (Fig. 3H), indicating a specific impact of SVCT2 overexpression in microglia on post-synaptic long-term plasticity.

Further investigation into basal synaptic transmission via input/output (I/O) curves recorded from Schaffer collateral-commissural fibers revealed augmented synaptic responsiveness in microglial SVCT2-injected mice compared to 5xFAD mice injected with control virus, as evidenced by a heightened maximal fEPSP slope (Fig. 3I), indicating that SVCT2-overexpressing 5xFAD mice present an enhanced synaptic efficacy upon robust stimulation.

Taken together, our electrophysiological data strongly suggest a pivotal role for microglia overexpressing SVCT2 in enhancing synaptic plasticity within the hippocampal circuitry of the 5xFAD mice. Hippocampal LTP was measured to assess synaptic plasticity, a physiological correlate of memory. The 5xFAD mice overexpressing SVCT2 in microglia showed a significantly higher LTP magnitude (Fig. 3D), indicating improved synaptic plasticity. Overall, the behavioral improvements observed in NOR and MWM tests and the increased synaptic plasticity underscore the beneficial impact of microglial SVCT2 overexpression during AD-like pathology progression in the 5xFAD mice.

### Overexpressing SVCT2 in Microglia Remodels the Synaptic Proteome Critical to Mitochondrial Energetics

To elucidate the proteomic underpinnings associated with SVCT2 overexpression in microglia in 5xFAD mice, we performed an in-depth high-throughput LC-MS/MS analysis targeting synaptic terminals (Fig. 4A). The results revealed a substantial reconfiguration of the synaptic proteome. Differential expression analysis indicated that 85 proteins were significantly upregulated and 52 were downregulated in the 5xFAD-SVCT2 group compared to the 5xFAD-Empty cohort (Fig. 4B). Subcellular localization analyses underscored a mitochondrial enrichment of altered proteins, suggesting the influence of microglial SVCT2 on synaptic mitochondrial proteomic composition and function (Fig. 4C).

**Figure 4:**
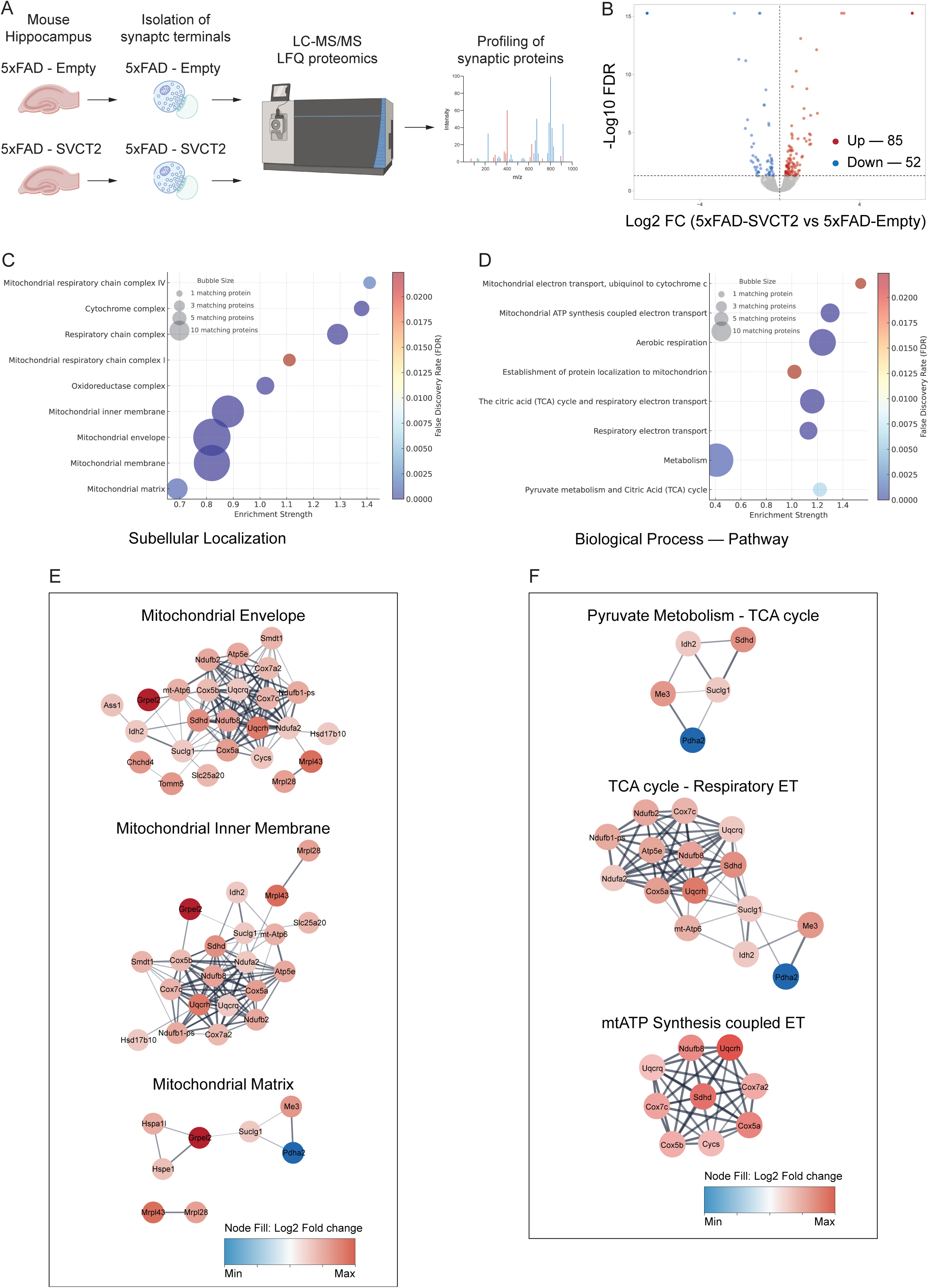
Reprogramming of 5xFAD synaptic mitochondrial proteome following microglial SVCT2 overexpression. A: Diagram illustrating the proteomic workflow, detailing the isolation of synaptic terminals from the hippocampi of 5xFAD mice, with and without SVCT2 overexpression, followed by LC-MS/MS analysis for quantitative proteomic profiling. B: Volcano plot showing differential protein expression between 5xFAD mice with SVCT2 overexpression (5xFAD-SVCT2) and controls (5xFAD-Empty), highlighting significantly upregulated (red) and downregulated (blue) proteins, with significance thresholds marked by dashed lines. Data are derived from 5 (5xFAD-SVCT2) mice and 4 (5xFAD-Empty) mice. C and D: Visualization of subcellular localization (C) and biological pathway (D) enrichment of differentially expressed proteins in 5xFAD-SVCT2 synapses. Bubble plots demonstrate the localization and functional categorization of proteins, with the size of each bubble representing the number of proteins and the color intensity reflecting the enrichment significance. E and F: Comprehensive network diagrams merging mitochondrial protein interactions (E) with detailed analyses of key metabolic pathways (F). Network diagrams highlight the interconnectedness of modulated proteins within the mitochondrial matrix and inner membrane and their involvement in pyruvate metabolism, the TCA cycle, and the respiratory electron transport chain.

Gene Ontology (GO) and pathway enrichment analyses have elucidated significant proteomic shifts, specifically highlighting an enhancement in the mitochondrial electron transport chain (ETC) and related components. Notably, the analysis revealed an upregulation in critical subunits of the mitochondrial respiratory chain Complex IV, such as Cox5a, Cox7a2, and Cox7c, suggesting a fortified mitochondrial respiratory function. This enhancement is complemented by an increase in ATP synthase component Atp5e, pointing to a potential uplift in ATP production capacity—a fundamental aspect for synaptic efficiency and adaptability (Fig. 4D).

Protein-protein interaction (PPI) analysis offers a detailed insight into mitochondrial protein dynamics, with alterations indicating an invigoration of mitochondrial functioning in response to SVCT2 overexpressing microglia (Fig. 4E and F). The elevation in the expression of Uqcrq and Uqcrh, components of the mitochondrial respiratory chain complex III, and Sdhd, a part of Complex II, mirrors a potential enhancement in the ETC’s efficiency facilitated by microglia overexpressing SVCT2 (Fig. 4E). The presence of proteins such as Ndufa2 and Ndufb8 from Complex I underscores a broad-based support for mitochondrial electron transport enhancement.

Within the mitochondrial matrix, the overexpression of SVCT2 in microglia markedly enhances the expression of pivotal proteins, thereby optimizing bioenergetic processes. The elevated levels of Hspa1l and Hspe1, chaperone proteins, enhance protein folding and stress response mechanisms, crucial for preserving mitochondrial functionality across different physiological states. The upregulation of Mrpl28, an integral component of the mitochondrial ribosome, signifies a bolstered synthesis of mitochondrial proteins, vital for the ETC and ATP production continuity. Furthermore, Grpel2’s role as a mitochondrial chaperone, involved in protein import and organizing the mitochondrial network, suggests an improved capacity for protein import and mitochondrial structural integrity. This is pivotal for the efficient operation of the mitochondria. Me3, which facilitates the malate/aspartate shuttle, is instrumental in efficiently transferring reducing equivalents into the mitochondria, enhancing NADH utilization for ATP synthesis. The observed increase in Suclg1, a crucial enzyme in the succinate-CoA ligase complex of the TCA cycle, highlights an enriched ability to synthesize ATP from TCA cycle intermediates. This enhancement underscores the critical function of the mitochondrial matrix in energy metabolism, pointing to a significant microglial overexpressing SVCT2-mediated boost in synaptic mitochondrial ATP production (Fig. 4E).

Further analysis of the interplay between pyruvate metabolism, the TCA cycle, the ETC, and ATP production delineates a suite of proteins pivotal to the synaptic mitochondria’s energy production pathways in 5xFAD mice, notably augmented by SVCT2 overexpression in microglia (Fig. 4F). The enhanced expression of Idh2 and Sdhd, crucial for the TCA cycle, marks a robust increase in NADH production. This uptick in NADH feeds into the ETC and likely amplifies synaptic ATP synthesis efficiency, further reflecting an optimized bioenergetic state. The upregulation of Complex III components, Uqcrq and Uqcrh, alongside Cycs—a critical player in the cytochrome c oxidase complex—illustrates a fortified electron transport mechanism. This enhancement ensures a seamless electron flow through the ETC, which is pivotal for sustaining high rates of oxidative phosphorylation and ATP generation. The involvement of Complex IV subunits, Cox5a, Cox7a2, and Cox7c, further solidifies the link between the TCA cycle and the ETC, facilitating efficient energy substrate conversion and bolstering mitochondrial respiratory function. Lastly, the increased levels of Atp5e and mt-Atp6, integral to the oxidative phosphorylation’s terminal phase, further signal a heightened ATP synthesis capability.

The enhanced expression profile of these mitochondrial proteins indicates a highly efficient and robust bioenergetic environment in the synaptic terminals of microglial SVCT2-overexpressing 5xFAD mice hippocampus. These findings provide a deeper understanding of the molecular changes through which microglial SVCT2 overexpression may ameliorate memory deficits and improve synaptic plasticity in the 5xFAD mice hippocampus.

### SVCT2 Overexpression Drives a Clearance Signature in Hippocampal 5xFAD Microglia

Our hypothesis posits that SVCT2 overexpression fosters an intracellular microenvironment that optimizes microglial functions for maintaining synaptic health. To unravel the molecular mechanisms through which SVCT2 overexpression exerts its effects, we performed a comprehensive transcriptomic analysis of microglia isolated from the hippocampus of 5xFAD mice.

RNA sequencing of MACS-isolated hippocampal 5xFAD microglia (Fig. 5A) revealed profound transcriptomic reprogramming upon SVCT2 overexpression compared to controls (5xFAD-Empty), as shown in Figure 5B. This extensive transcriptomic remodeling suggests a potential role for the SVCT2-dependent regulation of redox balance in orchestrating the functional state of 5xFAD microglia.

**Figure 5:**
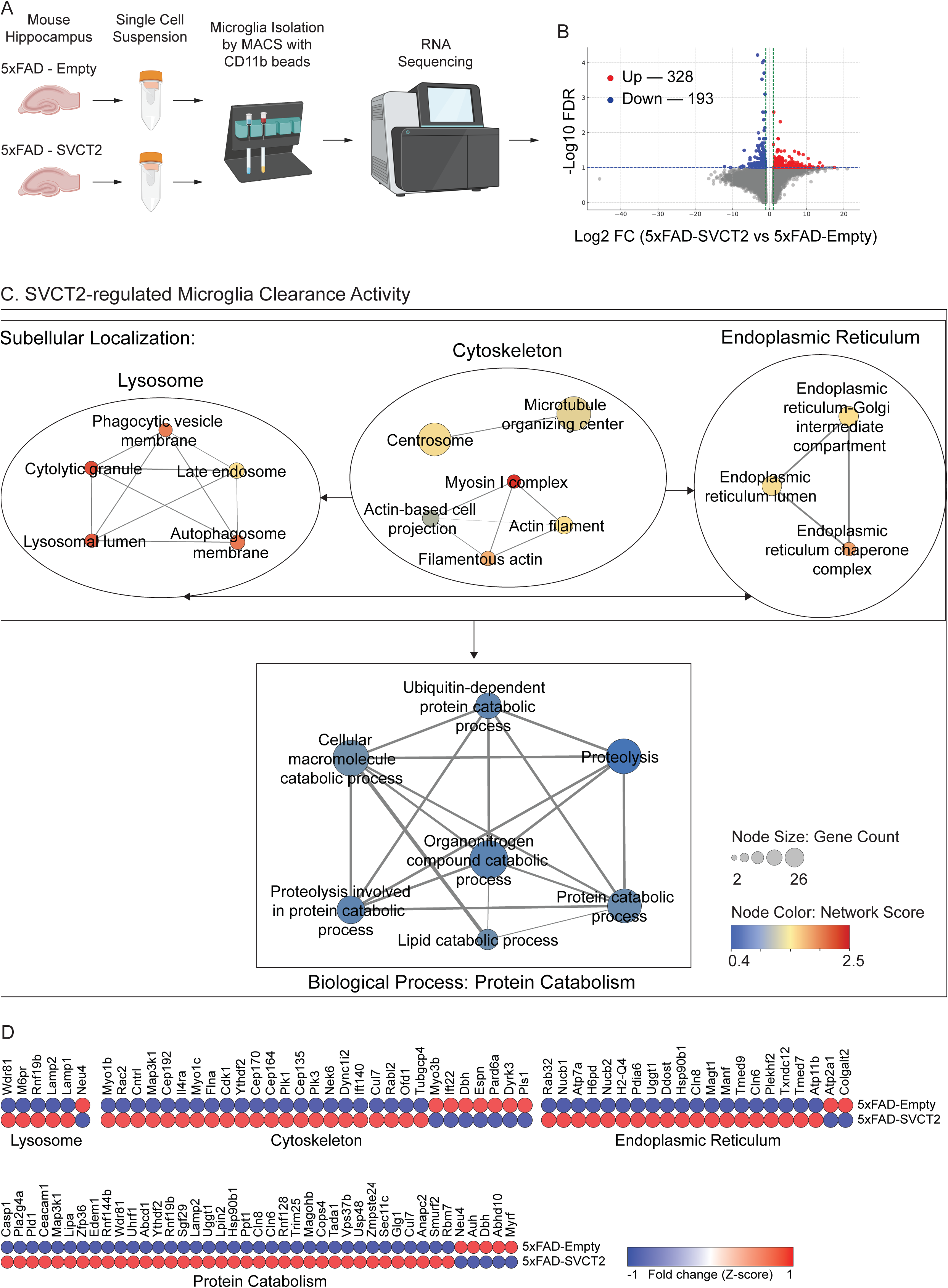
Clearance signature of SVCT2-overexpressing microglia in the hippocampus of 5xFAD mice. A: The workflow depicts the RNA sequencing process used to compare gene expression in microglia between 5xFAD-SVCT2 and 5xFAD-Empty (N=3). Hippocampal tissue was collected and dissociated into single cells, and microglia were isolated using CD11b+ magnetic-activated cell sorting. Post RNA extraction and cDNA library construction and sequencing were performed, with subsequent data analysis to identify differential expression patterns, providing insights into SVCT2’s role in microglial function. B: Volcano Plot of differentially expressed genes between 5xFAD-SVCT2 vs. 5xFAD-Empty conditions. Genes that are significantly upregulated are shown in red and downregulated in blue, with the x-axis representing the log2 fold change and the y-axis showing the - log10 of the false discovery rate (FDR). C: Network diagrams illustrating the subcellular localization and associated biological processes of the differentially expressed genes. It delineates the involvement of genes in lysosomal function, cytoskeletal organization, and the endoplasmic reticulum’s role in protein catabolism. D: Heatmap displaying gene expression changes across four main functional categories: lysosome, cytoskeleton, endoplasmic reticulum, and catabolism. Each row represents a gene, while columns are grouped by functional category. Gene expression levels are color-coded, with red indicating upregulation and blue indicating downregulation in 5xFAD-SVCT2 compared to 5xFAD-Empty.

Unbiased overrepresentation analysis *via* Gene Ontology fluxograms categorized the transcriptionally altered genes into clusters implicated in lysosomal function, cytoskeletal regulation, and endoplasmic reticulum (ER) stability (Fig. 5C). These changes are reflective of an augmented protein catabolism response (Fig. 5C), potentially amplifying the microglial clearance of amyloid deposits.

The differential gene expression heatmap (Fig. 5D) delineates significant transcriptional divergences in key lysosomal genes, highlighting the transcriptional landscape’s complexity. The pronounced upregulation of Lamp2, a critical component for maintaining lysosomal membrane integrity and facilitating autophagic processes, indicates an enhanced lysosomal degradation capability. This phenomenon is further supported by the increased transcription levels of M6pr, essential for the trafficking of hydrolases, suggesting a fortified enzymatic repertoire within the lysosomes. Concomitant transcriptional modifications in genes such as Wdr81 and Rnf19b unveil a sophisticated reorganization of lysosomal pathways, potentially augmenting the cell’s ability to process complex biomolecules. This may represent an adaptation to optimize cholesterol metabolism and glycosaminoglycan degradation.

In the cytoskeletal domain, the upregulation of Map3k1 in conjunction with Myo1b signifies a reinforcement of microtubule structures and actin filament dynamics, crucial for the precise regulation of microglial mobility and intracellular trafficking processes. The upregulation of Rac2 may indicate a strategic refinement of the actin cytoskeleton to optimize cellular mechanics and enhance phagocytic efficiency.

Within the endoplasmic reticulum, the transcriptional profile reveals an enhanced protein management system, as evidenced by the upregulation of Rab32. This increase may boost the endoplasmic reticulum’s capacity for protein folding and trafficking, reflecting a more dynamic and responsive protein quality control mechanism. The observed upregulation of Atp7a and H6pd could represent an adjustment of the endoplasmic reticulum stress responses, promoting a more stable and effective protein processing environment.

Regarding catabolic pathways, the upregulation of genes like Casp1 and Pla2g4a highlights an intensified protein turnover and degradation process, essential for maintaining cellular balance and preventing amyloid accumulation. The increased expression of Pld1 and Ceacam1 indicates a precise tuning of the ubiquitin-proteasome system, which is pivotal for preserving proteostasis.

Collectively, these transcriptional alterations delineate an advanced microglial functional framework steered by SVCT2 to optimize amyloid clearance and maintain microglial homeostasis.

### SVCT2 Overexpression Increases Microglia-Amyloid Interaction Enhancing Amyloid Clearance

The transcriptomic reconfiguration induced by SVCT2 overexpression in microglia markedly alters genes crucial for lysosomal function, cytoskeletal organization, and ER stability. This molecular shift implies an enhanced capacity of SVCT2-overexpressing microglia to clear amyloid deposits, suggesting a strategic approach to bolstering the brain’s defense against amyloid pathology.

Motivated by these findings, we investigated the effects of SVCT2 overexpression on the dynamics between microglia and amyloid in the hippocampus of 5xFAD mice. Our results revealed a substantial decrease in amyloid-beta accumulation in the hippocampi of 5xFAD-SVCT2 mice, as demonstrated by reduced BAM-10 stained area (Fig. 6A). Corresponding to this decrease in amyloid load, SVCT2 overexpression in 5xFAD microglia led to a notable shrinkage in amyloid plaque size and an enhancement in the extent of microglial clustering around amyloid plaques (Fig. 6B). These findings suggest a more efficient and focused microglial engagement with amyloid deposits following SVCT2 overexpression.

**Figure 6:**
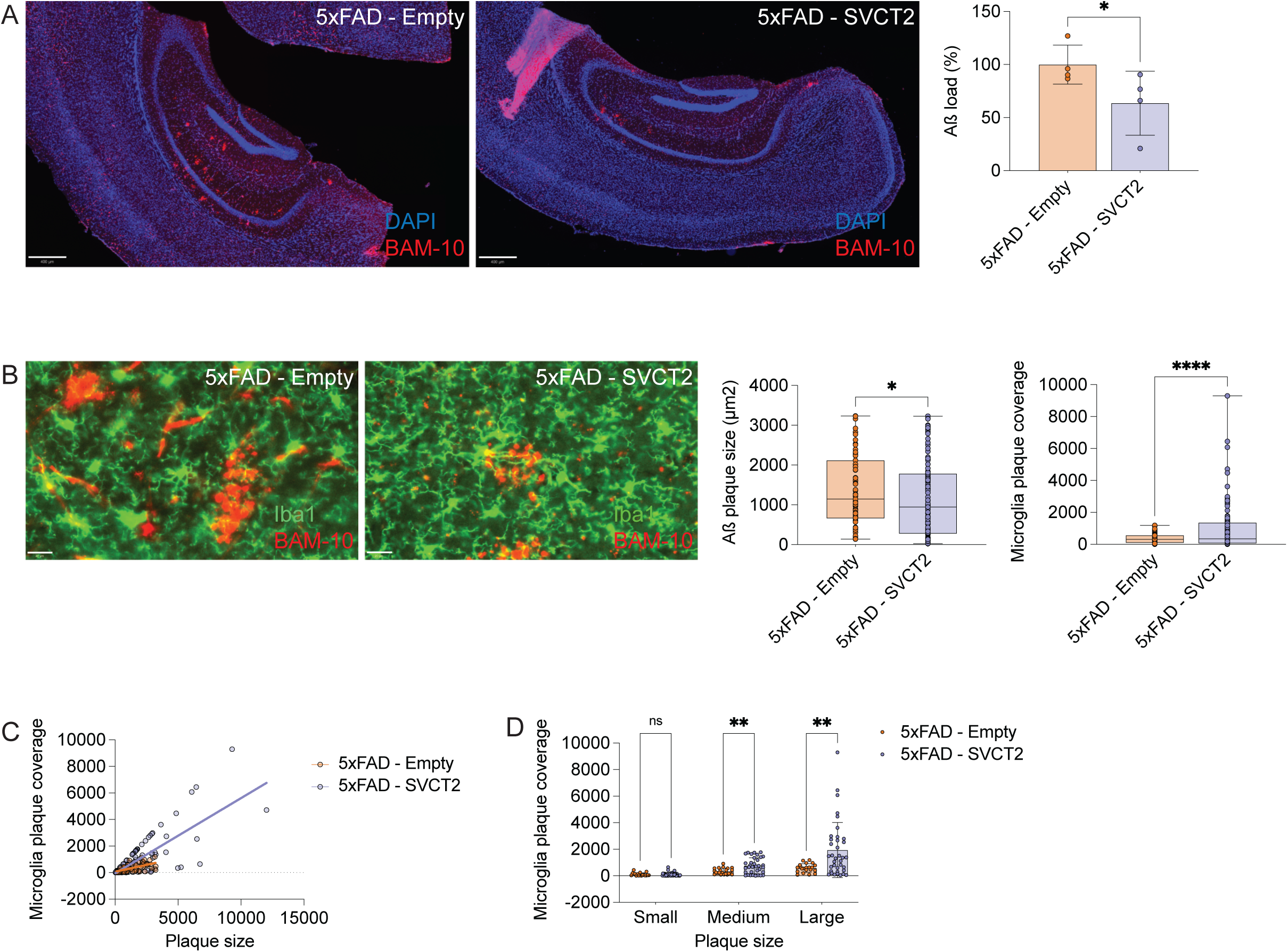
SVCT2 overexpression in microglia modulates amyloid plaque deposition in 5xFAD hippocampus. A: Fluorescence images displaying hippocampal amyloid plaques (labeled with BAM-10, red) in 5xFAD mice transduced with either Empty or SVCT2 constructs. Nuclei were stained with DAPI (blue). Scale bar: 400µm. The graph represents the quantification of amyloid plaque burden. Data are expressed as the mean ± SD of the BAM-10 stained area. Each data point in the graph represents an individual mouse. 5xFAD-Empty (N=4); 5xFAD-SVCT2 (N=4). * p<0.05 (Unpaired t-test). B: Fluorescence images displaying hippocampal amyloid plaques (labeled with BAM-10, red) and microglia (labeled with Iba1, green) in 5xFAD mice transduced with either Empty or SVCT2 constructs. Scale bar = 10µm. Graphs depict the quantification of amyloid plaque size (left) and microglial plaque coverage (right). Data represent 67 plaques from 4 different mice for 5xFAD - Empty and 102 plaques from 4 mice for 5xFAD - SVCT2, displayed as the mean ± SD. * p<0.05; **** p<0.0001 (Unpaired t Test). C: Regression analysis of microglial plaque coverage against amyloid plaque size in 5xFAD mice transduced with control and SVCT2 constructs. The scatter plot demonstrates a positive correlation between plaque size and microglial coverage in both groups. Data points represent individual plaques. D: Microglia clustering around different plaque size categories (Small, Medium, Large), derived from (B). The bar graph shows the mean microglial coverage for each plaque size category, displayed as mean ± SD, comparing 5xFAD mice transduced with Empty versus SVCT2 constructs. Data points represent individual plaques. **p<0.01 (Two-way ANOVA).

We conducted a regression analysis to solidify these observations, uncovering a significant positive association between plaque size and microglial encapsulation. For the 5xFAD-Empty mice, the analysis yielded a plaque size coefficient of 0.207 and a p-value of 1.71e-07, accounting for approximately 37.3% of the variance in microglial coverage of amyloid plaques (Fig. 6C). Conversely, the 5xFAD-SVCT2 group presented a plaque size coefficient of 0.566 and a p-value of 9.49e-22, revealing a more pronounced positive correlation and accounting for about 57.8% of the variance (Fig. 6C). These results points to SVCT2 overexpression fostering more substantial microglial aggregation around larger amyloid plaques.

Moreover, in comparison to the 5xFAD-Empty mice, the 5xFAD-SVCT2 group exhibited significantly increased microglial coverage of medium and large plaques, consistent with the regression analysis findings (Fig. 6D). For small plaques, coverage levels were comparable between the groups (Fig. 6D), emphasizing SVCT2 overexpression’s specific enhancement of microglial responses to larger amyloid accumulations. Together, these data on microglia-amyloid interactions support the RNAseq results and highlight an improved capacity for SVCT2-overexpressing microglia to clear amyloid deposits in the 5xFAD hippocampus.

### SVCT2 overexpression in APP/PS1 mice and its relevance for decreased amyloid load and excitatory synaptic content in the hippocampus

To reinforce our hypothesis regarding the neuroprotective role of microglial SVCT2 in AD-like pathology, we also carried out analyses in hippocampal microglia derived from APP/PS1 mice (Borchelt et al., 1997), another widely recognized Aß-depositing mice. Flow cytometry analysis revealed a diminished expression of SVCT2 in the microglia of 4-month-old APP/PS1 mice in comparison with WT littermates (Supplementary Figure 1A), as observed in 5xFAD mice, corroborating that SVCT2 downregulation in microglia occurs in both models. In line with our expectations, microglial overexpression of SVCT2 (confirmed in Supplementary Figure 1B) also decreased the APP/PS1 hippocampus amyloid burden (Supplementary Figures 1C and D). Furthermore, microglial SVCT2 overexpression also regulated the excitatory synapse content. We observed that APP/PS1 mice injected with the SVCT2 virus presented more excitatory synapses than APP/PS1 mice injected with the control vector (Supplementary Figure 1E).

## Discussion

Alzheimeŕs disease (AD) is a multifaceted and complex neurodegenerative disease that requires an in-depth examination of its molecular underpinnings. The 5xFAD mouse model, a critical AD research tool, facilitates this exploration, simulating some of the disease’s pathological and cognitive hallmarks and offering invaluable insights into molecular dynamics and progression patterns (Sanchez-Varo et al., 2022).

SVCT2, the ascorbate transporter in the CNS, is crucial for modulating microglial functioning (Portugal et al., 2017). We previously demonstrated that many neurodegenerative and neuroinflammatory stimuli, such as ischemia-reperfusion injury, uveitis, and LPS exposure, trigger the downregulation of this transporter in microglia (Portugal et al., 2017). With this in mind, we asked if SVCT2 downregulation in microglia also occurred in two Aß-depositing mice, the 5xFAD (Oakley et al., 2006) and the APP/PS1 (Borchelt et al., 1997), and whether SVCT2 overexpression in microglia could counteract AD-like pathology in both models. The importance of this question was reinforced by the significant downregulation of SVCT2 transcripts observed in human microglia associated with AD (Prater et al., 2023). In line with this, we observed a significant decrease in endogenous SVCT2 expression levels in microglia from both models. Of note, the decreased expression of SVCT2 in 5xFAD mice was observed during key stages of disease progression. This decrease, particularly noted between the 4^th^ and 6^th^ month, coincides with the rapid accumulation of amyloid-beta plaques and a marked decline in memory functions, implying a potential link between reduced SVCT2 expression, amyloid pathology, and the subsequent neurodegenerative cascade.

SVCT2 expression is crucial for modulating the redox balance in microglia (Portugal et al., 2017), which in turn could influence their phagocytic activity, inflammatory response, and overall capacity to manage amyloid plaques. The observed decrease in SVCT2 expression in 5xFAD could, therefore, be a contributing factor to the impaired clearance of amyloid-beta, exacerbating the pathological and cognitive symptoms characteristic of AD. Enhancing SVCT2 expression could bolster the microglial response to amyloid plaques, potentially reversing or mitigating the adverse effects of amyloid accumulation on neuronal and synaptic health. This hypothesis sets the stage for examining the broader implications of microglial SVCT2 overexpression on the neuronal and synaptic milieu.

Several mitochondrial proteins are altered in the AD brain (Minjarez et al., 2016). In line with this, we found a notorious reconfiguration of the synaptic mitochondrial proteome induced by SVCT2 overexpressing microglia in 5xFAD hippocampus, which likely plays a pivotal role in improving mitochondrial function related to increased synaptic plasticity and density (Turner et al., 2022). This proteomic remodeling, characterized by the upregulation of key mitochondrial ETC components and TCA cycle enzymes, potentially underpins an enhanced bioenergetic state conducive to neuronal and synaptic resilience (Tai et al., 2023). The upregulated ETC components maintain the mitochondrial membrane potential and ensure efficient ATP production (Nolfi-Donegan et al., 2020), the energy currency vital for synaptic plasticity. Furthermore, the increased expression of mitochondrial protein import machinery components hints at augmented mitochondrial biogenesis, suggesting an augmented capacity for assembling ETC complexes (Bogorodskiy et al., 2021). This, in turn, enhances the efficiency of oxidative phosphorylation, the chief process for energy production within the synapses. The elevation of TCA cycle enzymes further supports a revitalized metabolic flux, ensuring a steady supply of substrates for ATP generation (Tai et al., 2023), essential for sustaining synaptic activity and plasticity.

Mitochondrial dysfunction correlates with the accumulation of toxic amyloid species during the early stages of AD (Swerdlow et al., 2014). In line with this, the mitochondrial proteome reconfiguration not only signifies an optimized synaptic bioenergetic profile but also reflects a broader cellular strategy to counteract the AD-like neurodegenerative process in the 5xFAD hippocampus (Bogorodskiy et al., 2021; Swerdlow et al., 2014). The potential enhancements in mitochondrial function, brought about by the proteomic reconfiguration, might directly translate into notable improvements in synaptic plasticity and cognitive performance in the 5xFAD mice. The observed increase in LTP underscores the strengthened synaptic transmission and connectivity likely resulting from the mitochondrial enhancements driven by microglial SVCT2 overexpression. Thus, the intricate relationship between mitochondrial proteome remodeling and synaptic plasticity enhancements induced by SVCT2 overexpressing microglia counteract the bioenergetic deficits commonly associated with AD and potentially foster an environment that supports synaptic plasticity and function.

How does the overexpression of SVCT2 in microglia translate into such pronounced improvements in synaptic function? This line of inquiry directed our attention back to the microglial compartment. The hypothesis is that SVCT2 overexpression, by modulating the microglial response, plays a pivotal role in controlling the clearance activity of these cells. Our RNAseq analysis revealed significant transcriptional changes in SVCT2-overexpressing microglia within the 5xFAD hippocampus, particularly in cytoskeletal dynamics and lysosomal integrity genes. These molecular alterations indicate a profound reconfiguration of the microglial cytoskeletal framework, pivotal for the cells’ motility, phagocytic activity, and overall clearance capabilities. The upregulation of genes associated with microtubule stability and actin remodeling strongly suggests an enhancement in the microglia’s ability to rapidly mobilize towards and efficiently cluster around amyloid-beta plaques, a prerequisite for effective phagocytic engulfment and clearance of these pathological aggregates.

The reduction in amyloid burden due to enhanced microglial clearance has far-reaching implications for neuronal and synaptic health, particularly in the context of energy metabolism within the synapses (Wang et al., 2020). Amyloid plaques are known to disrupt synaptic function, partly by impairing mitochondrial dynamics and bioenergetics (Wang et al., 2020), leading to synaptic degeneration and cognitive decline. By mitigating the amyloid load, SVCT2-overexpressing microglia could indirectly contribute to preserving synaptic mitochondrial function and, consequently, the metabolic fitness of synapses.

The enhanced metabolic state promoted by SVCT2-overexpressing microglia can support many energy-intensive processes, such as the Na^+^/K^+^-ATP-ase and Ca^2+^ ATPases functioning, that are crucial to restore the neuronal membrane potential and neuronal excitability, and also regulating vesicle endo/exocytosis pathways, vital to glutamatergic receptor trafficking, regulating their bioavailability in the synapse, a pivotal step to LTP occurrence (Harris et al., 2012). These energy-intensive processes in the post-synapse are crucial for maintaining synaptic plasticity and overall cognitive function.

The intricate cascade from enhanced microglial clearance into improved cognitive outcomes involves a series of interconnected steps. Enhanced amyloid clearance relieves the synaptic mitochondria from amyloid-induced impairments, supporting efficient ATP production essential for synaptic transmission. This improved mitochondrial metabolic fitness at the synaptic level fosters synaptic plasticity, as evidenced by increased LTP, ultimately mitigating synaptic degeneration and memory deficits in the 5xFAD mice. Here, we revealed a complex and dynamic interaction between increased microglial SVCT2 expression, amyloid-beta accumulation, mitochondrial proteome remodeling, and synaptic plasticity in the context of AD-like pathology.

In this study, we elucidated the intricate and dynamic interplay between increased microglial SVCT2 expression, amyloid-beta accumulation, mitochondrial proteome alterations, and synaptic plasticity in the framework of AD-like pathology. Our data show a robust decrease of SVCT2 expression within microglia at crucial stages of AD-like pathology in the 5xFAD mice, a change that notably aligns with cognitive impairments. Remarkably, SVCT2 overexpression in microglia not only facilitated a significant reduction in amyloid plaque burden but also catalyzed a profound reshaping of the mitochondrial proteome within synapses, culminating in enhanced synaptic plasticity and improved memory performance. Overall, our data position microglial SVCT2 overexpression as a dual-action strategy that mitigates amyloid pathology and propels synaptic function for thwarting the neurodegenerative cascades characteristic of AD.

## Methods

### Experimental Models

Mice used in this work were bred at the i3S animal facility. Mice were housed under specific pathogen-free conditions in standard laboratory conditions with a light/dark cycle of 12 hours, 20°C, 45-55% humidity, and access to water and food *ad libitum*. All procedures were conducted according to the European Union guidelines for animal welfare (European Union Council Directive 2010/63/EU) and Portuguese law (DL 113/2013). All procedures considered Russell and Burch’s 3Rs principle. The Portuguese regulatory entity Direção Geral de Alimentação e Veterinária (DGAV) approved all experiments involving animal models (2022-02-18 003669). All efforts were made to minimize animal suffering and humane endpoints were adopted to safeguard animal welfare.

#### 5xFAD Mouse Model

The 5xFAD transgenic mouse model overexpresses five familial Alzheimer’s disease (FAD) mutations in human APP and PS1 [APP: K670N/M671L (Swedish) + I716V (Florida) + V717I (London); PS1: M146L + L286V] (Oakley et al., 2006) Both transgenes are under the regulation of the *Thy1* promoter. Founders were purchased from The Jackson Laboratory (MMRRC stock #34840). This mouse line was rederived and maintained in a C57BL/6J background. The breeding scheme implemented was informed by the observation that female 5xFAD subjects exhibit a more rapid disease progression (Forner et al., 2021). As such, we selectively bred males heterozygous for the 5xFAD transgene with wild-type females. Progeny was genotyped using the following primers: Primer mutant (5’ to 3’): AAG CTA GCT GCA GTA ACG CCA TTT; Primer WT (5’ to 3’): ACC TGC ATG TGA ACC CAG TAT TCT ATC; Primer common (5’ to 3’): CTA CAG CCC CTC TCC AAG GTT TAT AG. This study exclusively utilized female 5xFAD mice with heterozygous genetic profiles.

#### AßPPswe/PS1A246E Mouse Model

The AßPPswe/PS1A246E, hereafter, APP/PS1 mice, co-expresses a chimeric mouse-human amyloid-ß protein precursor (AßPP) bearing a human Aß domain with mutations (K595N and M596L) linked to Swedish familial AD pedigrees and human presenilin-1 A246E mutation, with both transgenes under the control of the mouse prion protein promoter (Borchelt et al., 1997). APP/PS1 mice were genotyped by PCR using PSEN primers (5’ to 3’): AAT AGA GAA CGG CAG GAG CA (forward) and GCC ATG AGG GCA CTA ATC AT (reverse); APP primers (5’ to 3’): GAC TGA CCA CTC GAC CAG GTT CTG (forward) and CTT GTA AGT TGG ATT CTC ATA TCC G (reverse); WT primers (5’ to 3’): CCT CTT TGT GAC TAT GGT GAC TGA TGT CGG (forward) and GTG GAT AAC CCC TCC CCC AGC CTA GAC C (reverse). APP/PS1 mice were maintained in SV129 background. This study exclusively utilized female APP/PS1 mice with heterozygous genetic profiles.

### Experimental Procedures

#### Behavioral tests

Procedures were conducted in the dark phase of the light/dark cycle and performed blind to genotypes. The tests were carried out in the following order: (1) Novel Object Recognition followed by an interval of 6 days, and (2) Morris Water Maze. The timeline is displayed in Figure 3A.

##### Novel Object Recognition (NOR)

The NOR test is employed to evaluate recognition memory in mice as before (Socodato et al., 2023; Socodato, Portugal, et al., 2020). This test is divided into three parts. In the first phase, the habituation, mice were placed in a box (40cm x 40cm x 40cm) and allowed to explore the apparatus for 10 minutes (Fig. 3B). After 24 hours, mice were put in the same box in the presence of two identical objects equidistant from each other (blue squares; Figure 3B). Free exploration in the presence of both objects was allowed for 10 minutes. Four hours later, one of the familiar objects was replaced by a new one that differed from the previous one in color, shape, and texture (yellow triangle; Fig. 3B). Mice were again positioned in the apparatus, and 3 minutes were given for assessment. Exploration was defined as follows: the mouse touched the object with its nose, or its nose was directed toward the object at a distance shorter than 2cm. The analyses for the test day were performed using the Boris software (Friard & Gamba, 2016) to extract the exploration time of the novel (Time_Novel_) and familiar (Time_Familiar_) objects. After that, the percentage of recognition was calculated using the following ratio:

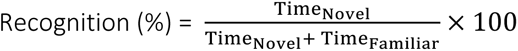

##### Morris Water Maze (MWM)

The MWM was used to evaluate spatial memory. The apparatus consisted of a circular pool (110 cm diameter) of fiberglass filled with water (21±1°C). This test is divided into three consecutive steps. Firstly, the cued training was conducted for two days with four trials *per* animal *per* day. A non-visible escape platform (7 x 8 cm) was submerged 1 cm below the water surface in the quadrant center. Mice were trained to find the platform with a visual clue. Animals were subjected to four swimming trials with different start and goal positions. The training phase occurs in five days, with four trials *per* animal *per* day and 1 minute for accomplishment. The visual clue on top of the platform was removed, but others were placed across the room walls to identify the platform position. Only the mice are released from different positions in this stage, while the platform’s location remains unaltered. Once again, if the mice could not reach the platform in 1 minute, they were guided to the platform.

Nonetheless, 10 seconds would be given to the mice on top of the platform for each trial, allowing them to learn its location. The test day occurs 24 hours after the last training. In this stage, the platform was removed, challenging the mice to recall where the platform was supposed to be. Each mouse was released from the same position (opposite quadrant) and allowed to freely swim around the apparatus for 30 seconds to search the platform’s position. All parameters were automatically evaluated by SMART v3.0 software (Panlab, Barcelona, Spain).

### Stereotaxic injection of Adeno-Associated Virus (AAV) into the hippocampus

The mice were initially sedated using a 3% isoflurane anesthesia, followed by the subcutaneous administration of the analgesic buprenorphine at a dosage of 0.08 mg/kg to ensure adequate pain management throughout the procedure. Subsequently, each mouse was carefully positioned within a digitally controlled stereotaxic apparatus equipped with an integrated microinjection system (Stoelting Co.). To maintain the depth of anesthesia while minimizing respiratory depression, the isoflurane concentration was adjusted to 1%. A rectal thermal probe linked to a rodent warming unit (Stoelting Co.) was employed to sustain the subject’s body temperature at a physiological norm of 37°C. Precise hippocampal targeting was achieved by using the bregma as a landmark, with the following coordinates: Anteroposterior (AP) at -2.0 mm; Mediolateral (ML) at ±1.3 mm; and Dorsoventral (DV) at -2.2 mm.

Afterward, stereotaxic delivery of custom-engineered AAV9 vectors was carried out. These vectors harbored a bicistronic transgene payload, encoding for SVCT2 and mCherry proteins (the “SVCT2 virus”). An alternative construct containing only the mCherry reporter (the “empty virus”) was a control. The expression of these transgenes was driven by the minimal CD68 promoter, chosen for its specificity to the target microglia within the hippocampal region. Both 5xFAD transgenic mice and their wild-type littermates were administered the viral vectors at the age of two months. The impact and efficacy of transgene delivery were subsequently assessed four months post-administration when the mice reached six months of age.

### Tissue dissociation and microglial enrichment

#### Percoll Gradient

As outlined in previous studies, the method for separating cells employed a 24% isotonic Percoll gradient (Galatro et al., 2017). In brief, mice were euthanized under deep anesthesia induced by pentobarbital (400 mg/kg) and subsequently perfused with ice-cold phosphate-buffered saline (PBS). The brains were carefully extracted and subjected to mechanical dissociation using a glass tissue grinder in Hank’s Balanced Salt Solution (HBSS), enhanced with 15 mM HEPES and 0.6% glucose. The resulting tissue homogenates were passed through 70 μm cell strainers that had been moistened in advance to achieve single-cell suspensions. These suspensions were centrifuged at 300 rcf for 10 minutes at a temperature of 4°C. After discarding the supernatant, the cell pellets were resuspended in an isotonic solution of 24% Percoll, over which PBS was gently layered. The samples were then centrifuged at 800 rcf for 30 minutes at 4°C to form gradients. After centrifugation, the cell pellets were again suspended in HBSS containing 15 mM HEPES, 0.6% glucose, and 1 mM EDTA, followed by two washing steps. The purified cells were finally readied for analysis via flow cytometry.

#### Microglia Isolation Using MACS

To isolate microglia from the hippocampus, mice were anesthetized with pentobarbital (400 mg/kg) and perfused with ice-cold PBS. The hippocampi were dissected and then dissociated mechanically in ice-cold Dounce buffer, which contained 15 mM HEPES, 0.5% glucose, 160 U/mL DNase, and 20 U/mL RNasin® Plus Ribonuclease Inhibitor, adjusted to pH 7.4. The resulting cell suspensions were centrifuged at 300 rcf for 10 minutes at 4°C. The cell pellets were resuspended in MACS buffer (0.5% BSA and 1 mM EDTA in PBS, pH 7.4) and incubated with CD11b MicroBeads (Miltenyi Biotec) for 30 minutes to label the microglial cells. After removing excess beads through centrifugation, the cells were passed through LS columns (Miltenyi Biotec) mounted on a quadroMACS separator (Miltenyi Biotec) and pre-rinsed with MACS buffer. The columns were washed, and the CD11b-positive cells were eluted, collected by centrifugation, and set aside for either flow cytometry analysis or further processing for RNA isolation and sequencing.

Microglia isolated for RNA sequencing underwent a second round of purification by being passed through a new LS column to enhance purity further. After washing, the enriched CD11b-positive cells were lysed using RLT buffer plus (Qiagen) supplemented with 40 μM DTT. RNA was then isolated from these lysates using the Qiagen RNeasy kit, following the manufacturer’s protocol. The Bioanalyzer 2100 RNA Pico chips assessed the RNA integrity according to the manufacturer’s protocol.

### Flow cytometry

#### Analysis of whole brain microglia

Following the tissue dissociation procedure outlined above using a 24% Percoll gradient, the resultant cell suspensions were quantified using the Countess™ Automated Cell Counter (Invitrogen). A total of 2.5 × 10^6 viable cells were allocated to each well of a U-bottom 96-well plate. The cells were first stained with Zombie Green™ viability dye (diluted 1:500 in PBS) for 30 minutes to differentiate live cells. After Fc receptor blocking using CD16/CD32 antibodies (1:100 dilution), the cells were labeled with CD45 (PE-conjugated, 1:200 dilution) and CD11b (PE/Cy7 conjugated, 1:200 dilution) markers. Post-staining, cells were fixed with 2% paraformaldehyde (PFA) and permeabilized using eBioscience™ Permeabilization Buffer to allow intracellular staining. Intracellular SVCT2 was detected using a primary antibody (1:100 dilution) followed by incubation with Alexa Fluor™ 647-conjugated goat anti-rabbit IgG secondary antibody (1:500 dilution). After washing, samples were analyzed on an LSRFortessa flow cytometer (BD Biosciences). Compensation was automatically set using monolabel controls (cells stained with individual antibodies or dyes), and SVCT2 staining was validated using a fluorescence-minus-one (FMO) control. Data analysis was conducted using FlowJo X10 Software (TreeStar).

#### Analysis of hippocampal microglia

Microglia isolated from the hippocampus, as described in the previous section using the MACS system, were prepared similarly. After plating and PBS washes, cells were stained with Zombie Green™ (1:500 in PBS) for viability assessment. Fc receptors were blocked (CD16/CD32, 1:100 dilution) before staining with CD11b-PE/Cy7. Following fixation and permeabilization (2% PFA and eBioscience™ Permeabilization Buffer, respectively), cells were incubated with anti-SVCT2 and anti-mCherry antibodies (both 1:100). Secondary antibodies, Alexa Fluor™ 568 for mCherry and Alexa Fluor™ 647 for SVCT2 (both 1:500), were then applied. After a final wash, cells were analyzed using the LSRFortessa. Compensation and FMO controls were set up as described above, including specific FMO controls for SVCT2 and mCherry. Data analysis was performed using FlowJo X10 Software.

### Tissue preparation for immunofluorescence analysis

Mice were euthanized using a pentobarbital dose of 400 mg/kg and subsequently perfused with ice-cold PBS to clear the blood. The brains were carefully excised and fixed in 4% paraformaldehyde (PFA) for a full day to ensure thorough fixation. Following fixation, the brains were rinsed with PBS and then subjected to cryoprotection. This was achieved by immersing the tissue in successive solutions of 15% followed by 30% sucrose, each supplemented with 0.01% sodium azide to prevent microbial growth. For sectioning, the cryoprotected brains were embedded in an Optimal Cutting Temperature (OCT) compound to support the tissue structure. They were then rapidly frozen to preserve cellular integrity and sectioned at a thickness of 30 µm using a Leica CM3050S Cryostat, ensuring consistent slice thickness for uniform staining. The brain sections underwent a permeabilization step with 1% Triton X-100 for 15 minutes to increase antibody accessibility. This was followed by a quenching step with 0.2M NH4Cl for another 15 minutes to reduce autofluorescence. For one hour, sections were incubated in a 1% BSA solution within the permeabilization buffer to block non-specific binding. Primary antibodies targeting Iba-1, mCherry, and BAM-10 (diluted at 1:400, 1:400, and 1:100, respectively, in blocking solution) were applied to the sections, which were then incubated at 4°C for 72 hours to allow for thorough binding. After primary antibody incubation, sections were washed with PBS and incubated with fluorescently labeled secondary antibodies (Alexa Fluor™ 647, Alexa Fluor™ 488, and Alexa Fluor™ 568 conjugated antibodies, all at 1:1000 dilution) in blocking solution for 24 hours at 4°C. Following secondary antibody incubation, sections were washed and stained with DAPI (125 µg/mL) for 15 minutes to label nuclei, followed by a final PBS wash. The staining procedures were conducted with gentle orbital agitation to ensure even staining. After the final wash, brain sections were mounted onto glass slides. Excess PBS was carefully removed, and a 90% glycerol was applied. One hour later, the slides were sealed with nail polish to prevent drying and left to cure in a dark environment for at least 24 hours before imaging.

### Imaging acquisition

#### High-Throughput Imaging Analysis of Amyloid and Microglia

To elucidate the dynamic interplay between amyloid beta (Aβ) deposition (BAM-10 labeling) and microglia (Iba1 labeling) within the 5xFAD model hippocampus, our methodology leveraged the PhenoImager HT system for high-throughput imaging of tissue slides. This system’s automation and rapid scanning capabilities were crucial, efficiently acquiring comprehensive whole-slide images within a 15mm x 15mm area. In the post-imaging phase, we employed QuPath software for the rigorous quantification and characterization of Aβ deposition and microglial distribution patterns.

#### Confocal imaging

High-resolution images of tissue sections from both the neocortex and dorsal hippocampus were captured using a Leica TCS SP5 confocal microscope. The imaging was conducted in 8-bit sequential mode at a scan speed of 400 Hz, utilizing the standard TCS mode to ensure optimal resolution and minimize signal noise. The pinhole was adjusted to one Airy unit across all imaging sessions to achieve consistent optical sectioning. Images were generated using various laser excitation combinations to illuminate the samples effectively. The resolution of the acquired images was set to either 512 x 512 or 1024 x 1024 pixels, depending on the detail required for the analysis. Hybrid Detector (HyD) technology was employed for its superior sensitivity and dynamic range, facilitating the capture of high-quality images. Comprehensive Z-stacks were collected for each tissue section to encompass the entire sample depth, ensuring a thorough volumetric analysis. To maintain consistency and allow accurate sample comparison, equivalent stereological regions were delineated and imaged across all tissue sections within a given slide. This approach ensured that data collection was systematic and representative of the entire tissue architecture.

### Electrophysiology

#### Preparation of Acute Hippocampal Slices

Acute hippocampal slices were meticulously prepared precisely as before (Rei et al., 2020). 5xFAD mice and WT littermates were injected for AAV delivery as described above. Mice aged between 6- and 6.5-months were humanely euthanized using a rapid cervical dislocation followed by decapitation. Brains were swiftly excised and submerged in ice-cold artificial cerebrospinal fluid (aCSF) containing (in mM): 124 NaCl, 3 KCl, 1.2 NaH_2_PO_4_, 25 NaHCO_3_, 2 CaCl_2_, 1 MgSO_4,_ and 10 glucose, continuously oxygenated with a 95% O_2_ and 5% CO2 mixture. Hippocampi were carefully dissected and sectioned perpendicularly along their long axis into 400 μm thick slices using a precision tissue chopper. These slices were then incubated in oxygenated aCSF at room temperature (22–25°C) for at least 1 hour to ensure functional and energetic recovery before electrophysiological assessments.

#### Extracellular fEPSP Recordings

Post-recovery, slices were positioned in a specialized recording chamber designed for submerged slices, with a constant superfusion of warm (32°C) oxygenated aCSF at a 3 ml/min flow rate. Extracellular fEPSPs were recorded in the CA1 stratum radiatum using microelectrodes (4-8 MΩ) filled with aCSF. Stimuli were delivered to Schaffer collateral fibers *via* a bipolar concentric electrode (platinum/iridium, 25 μm diameter, <800 kΩ impedance), with each stimulus comprising a 0.1 ms pulse at 20-second intervals. Data acquisition was managed with an Axoclamp-2B amplifier and analyzed using WinLTP software (Anderson & Collingridge, 2001). The initial phase slope of averaged fEPSPs (from six consecutive responses) was quantified, setting the stimulus intensity to achieve a near 0.5 mV/ms slope, indicative of approximately 50% maximal response. Input/output (I/O) relationships were delineated by incrementally increasing stimulus intensity (20µA every 4 minutes, ranging from 60 to 340 µA), plotting fEPSP slopes against stimulus intensities. Maximal slope values were extrapolated from non-linear I/O curve fitting, utilizing an F-test for parameter differentiation. Paired-pulse facilitation (PPF) was evaluated by the slope ratio of two consecutive fEPSPs (fEPSP1/fEPSP0) with a 50ms interstimulus gap. Six paired responses were averaged for each PPF metric.

Long-term potentiation (LTP) was induced using a θ-burst stimulation pattern, which is more physiologically relevant to learning and memory processes (Albensi et al., 2007). This involved a single train of four bursts (200ms apart), each containing four 100Hz pulses. LTP magnitude was gauged as the percentage change in fEPSP slope (50-60 minutes post-induction) relative to pre-induction averages (10 minutes prior). Post-tetanic potentiation (PTP), reflecting transient transmitter release enhancements due to residual Ca^2+^ during high-frequency activity (Korogod et al., 2007), was assessed by averaging fEPSP slopes in the initial 4 minutes following LTP induction (Habets & Borst, 2007). Recording protocols were systematized to capture PPF data via one pathway stimulation, I/O metrics through an alternate pathway, and PTP and LTP measurements by reverting to the initial pathway. Baseline fEPSP stability was confirmed under standard stimulation conditions for over 10 minutes before any protocol adjustments. Each slice, representing two independent pathways, was utilized for one set of experiments per day.

### Enzyme-linked immunosorbent assay (ELISA) for Aß_1-42_ detection in soluble and insoluble fractions

Hippocampi were carefully dissected, immediately snap-frozen in dry ice, and stored at -80°C until further processing. Tissue was then thawed and homogenized in 200µL of ice-cold homogenization buffer (50mM Tris buffer, 4mM EDTA, 0.2% Triton X-100, pH 7.4 supplemented with a protease inhibitor cocktail). The homogenates were centrifuged, and the supernatant was saved as a soluble fraction. The pellets were then homogenized in 70% Formic acid and neutralized with 1mM Tris, pH11, the neutralized insoluble fraction. Both fractions were then used to measure the amyloid beta 1-42 (Aβ1-42) using a specific ELISA kit (Invitrogen) following the manufacturer’s instructions. Absorbance was measured by a multimode microplate reader (Synergy HT, Biotek, USA). The Aβ_1-42_ concentration was calculated by extrapolating from a standard concentration curve.

### Regression analysis for microglial SVCT2, spatial memory, and age

The relationship between SVCT2 expression, spatial memory performance, and age was investigated in a cohort of 5XFAD mice, focusing on two pivotal age milestones— 4 and 6 months—to capture the progression of AD-like pathology. SVCT2 expression levels were quantified, and spatial memory was assessed using the Morris Water Maze (MWM) test. Linear regression analyses were employed to examine the impact of age on SVCT2 expression and spatial memory performance. Age was the independent variable, while SVCT2 expression levels and spatial memory scores were the dependent variables. The analyses were designed to accommodate any potential non-linear relationships and variance inconsistencies within the data. Model efficacy was evaluated by examining R-squared values and standard error metrics. Statistical significance was determined with a p-value cutoff of <0.05. Additionally, 95% confidence intervals were calculated for the regression estimates. The regression modeling and subsequent analyses were conducted using Python (Version 3.8), specifically using the statsmodels library for the regression analysis and matplotlib and seaborn libraries for plot representation.

### Library preparation, RNA sequencing, and bioinformatics

Ion Torrent sequencing libraries were prepared according to the AmpliSeq Library prep kit protocol as we did before (Canedo et al., 2021; Socodato et al., 2023). Briefly, 1 ng of highly intact total RNA was reverse transcribed. The resulting cDNA was amplified for 16 cycles by adding PCR Master Mix and the AmpliSeq mouse transcriptome gene expression primer pool. Amplicons were digested with the proprietary FuPa enzyme, and then barcoded adapters were ligated onto the target amplicons. The library amplicons were bound to magnetic beads, and residual reaction components were washed off.

Libraries were amplified, re-purified, and individually quantified using Agilent TapeStation High Sensitivity tape. Individual libraries were diluted to a 50 pM concentration and pooled equally. Emulsion PCR, templating, and 550 chip loading were performed with an Ion Chef Instrument (Thermo Scientific MA, USA). Sequencing was performed on an Ion S5XL™ sequencer (Thermo Scientific MA, USA) as we did before (Canedo et al., 2021).

Data from the S5 XL run was processed using the Ion Torrent platform-specific pipeline software Torrent Suite v5.12 to generate sequence reads, trim adapter sequences, filter and remove poor signal reads, and split the reads according to the barcode. FASTQ and BAM files were generated using the Torrent Suite plugin FileExporter v5.12. Automated data analysis was done with Torrent Suite™ Software using the Ion AmpliSeq™ RNA plugin v.5.12 and target region AmpliSeq_Mouse_Transcriptome_V1_Designed as we did before (Canedo et al., 2021).

Raw data was loaded into Transcriptome Analysis Console (4.0 Thermo Fisher Scientific, MA, EUA) and first filtered based on ANOVA eBayes using the Limma package and displayed as fold change. Significant changes had a fold change of <-1.5 and >1.5, p-value < 0.05 and FDR < 0.1. Functional enrichment analyses were performed using STRING (Szklarczyk et al., 2019). Pathway enrichment was conducted using the REACTOME database with default settings. Enrichment scores for gene sets were calculated using an FDR cutoff of 0.05. Enriched pathways were manually recategorized to core transcriptomic modules and displayed as a network (constructed using Cytoscape).

### Synaptosomal preparations

Synaptosomes were acutely prepared as before (Costa-Pinto et al., 2024; Socodato et al., 2023; Socodato, Henriques, et al., 2020). One hundred micrograms of synaptosomal proteins from each sample were processed for proteomic analysis following the solid-phase-enhanced sample-preparation protocol as before (Osório et al., 2021). Enzymatic digestion was performed with trypsin/LysC (2 micrograms) overnight at 37◦C at 1000 rpm. The resulting peptide concentration was measured by fluorescence.

### High-throughput proteomics, data acquisition, and quantification

NanoLC-MS/MS performed protein identification and quantitation as before (Costa-Pinto et al., 2024; Socodato et al., 2023). This equipment comprises an Ultimate 3000 liquid chromatography system coupled to a Q-Exactive Hybrid Quadrupole-Orbitrap mass spectrometer (Thermo Scientific). Five hundred nanograms of peptides of each sample were loaded onto a trapping cartridge in a mobile phase of 2% ACN, 0.1% FA at 10μL per minute. After 3 minutes of loading, the trap column was switched in-line to a 50cm x 75μm inner diameter EASY-Spray column at 250nL per minute. Separation was achieved by mixing A (0.1% FA) and B (80% ACN, 0.1% FA) with the following gradient: 5 minutes (2.5% B to 10% B), 120 minutes (10% B to 30% B), 20 minutes (30% B to 50% B), 5 minutes (50% B to 99% B), and 10 minutes (hold 99% B). Afterward, the column was equilibrated with 2.5% B for 17 minutes. Data acquisition was controlled by Xcalibur 4.0 and Tune 2.11 software (Thermo Scientific).

The mass spectrometer was operated in the data-dependent (DD) positive acquisition mode alternating between a full scan (m/z 380-1580) and subsequent HCD MS/MS of the 10 most intense peaks from a full scan (normalized collision energy of 27%). The ESI spray voltage was 1.9kV. The global settings were: use lock masses best (m/z 445.12003), lock mass injection Full MS and chrom. peak width (FWHM) of 15 seconds. The full scan settings were as follows: 70k resolution (m/z 200), AGC target 3 x 106, maximum injection time 120 milliseconds; DD settings: minimum AGC target 8 x 103, intensity threshold 7.3 x 104, charge exclusion: unassigned, 1, 8, >8, peptide match preferred, exclude isotopes on, and dynamic exclusion 45 seconds. The MS2 settings were as follows: micro scans 1, resolution 35k (m/z 200), AGC target 2 x 105, maximum injection time 110 milliseconds, isolation window 2.0m/z, isolation offset 0.0m/z, dynamic first mass, and spectrum data type profile.

The raw data was processed using the Proteome Discoverer 2.5.0.400 software (Thermo Scientific) and searched against the UniProt database for the reviewed Mus musculus Proteome (2021_03 with 17,077 entries). A common protein contaminant list from MaxQuant was also considered in the analysis. The Sequest HT search engine was used to identify tryptic peptides. The ion mass tolerance was 10ppm for precursor ions and 0.02Da for fragment ions. The maximum allowed missing cleavage sites was set to two. Cysteine carbamidomethylation was defined as a constant modification. Methionine oxidation, deamidation of glutamine and asparagine, peptide terminus glutamine to pyroglutamate, and protein N-terminus acetylation, Met-loss, and Met-loss+acetyl were defined as variable modifications. Peptide confidence was set to high. The processing node Percolator was enabled with the following settings: maximum delta Cn 0.05; decoy database search target false discovery rate 1%, validation based on q-value. Protein label-free quantitation was performed with the Minora feature detector node at the processing step. Precursor ions quantification was conducted at the consensus step with the following parameters: unique plus razor peptides were considered, precursor abundance was based on intensity, and normalization was based on total peptide amount.

For the determination of DE between groups, the following filters were used: (1) only master proteins detected with high/medium confidence FDR; (2) a protein/phosphoprotein must be detected in more than 50% of samples in each experimental group (except for proteins that were depleted entirely in one of the experimental groups); (3) the *p*-value adjusted using Benjamini–Hochberg correction for the FDR was set to ≤ 0.05; (4) at least 50% of samples with protein-related peptides sequenced by MS/MS.

### Proteomics network analyses

For constructing granular networks, DE proteins retrieved from the LFQ experience were uploaded to STRING (Szklarczyk et al., 2019) within Cytoscape as before (Socodato et al., 2023). Enrichment analyses (FDR cutoff < 0.05 with the Benjamini-Hochberg multiple test adjustments) were carried out with REACTOME and GO as functional databases. Network construction and topography were carried out following manual pathway annotation and clustering in Cytoscape.

### Quantification and Statistical Analysis

Experimenters were blinded to genotypes and housing conditions. Data were tested for Gaussian distribution using the D’Agostino-Pearson omnibus test. All tests considered mice or specific cells as experimental units with a significance level of P < 0.05. Descriptive statistics, including mean, median, standard deviation, and interquartile range, were provided for each dataset and detailed within the Figure Legends accompanying the results. All statistical analyses and the creation of graphical representations of the data were conducted using GraphPad Prism (version 9.0.2 for macOS) and Python, ensuring rigorous and accurate data analysis and visualization.

The assembly of figure panels for publication was performed using Adobe Illustrator 2020 (version 24.3). Schematic diagrams were created using BioRender.

## Funding

This work was funded by Portuguese funds through FCT - Fundação para a Ciência e a Tecnologia under framework of the project EXPL/MED-NEU/0588/2021.

Fundação para a Ciência e a Tecnologia (MCTES) through Fundos do Orçamento de Estado also supported the work realized on Instituto de Medicina Molecular João Lobo Antunes by Sandra H. Vaz and Sara Costa-Pinto under the project PTDC/BTM-SAL/32147/2017.

2021 International Society for Neurochemistry (ISN) Career Development Grant funded the work realized on Instituto de Medicina Molecular João Lobo Antunes by Sandra H. Vaz

H2020-WIDESPREAD-05-2017-Twinning (EpiEpinet) under grant agreement No. 952455 funded the work realized on Instituto de Medicina Molecular João Lobo Antunes by Sandra H. Vaz and Sara Costa-Pinto.

This work was funded (in part) by Programa Operacional Regional do Norte and co-funded by European Regional Development Fund under the project “The Porto Comprehensive Cancer Center” with the reference NORTE-01-0145-FEDER-072678 - Consórcio PORTO.CCC – Porto.Comprehensive Cancer Center.

Camila C. Portugal and Renato Socodato hold an employment contract financed by national funds through FCT - in the context of the program-contract described in paragraphs 4, 5, and 6 of art. 23 of Law no. 57/2016, of August 29, as amended by Law no. 57/2017 of July 2019 (DL 57/2016/CP1355/CT0025 for Camila C. Portugal and DL 57/2016/CP1355/CT0024 for Renato Socodato).

Sara Costa-Pinto, Tiago O. Almeida, and Joana Tedim-Moreira hold Ph.D. fellowships funded by FCT (SFRH/BD/147277/2019, SFRH/BD/147981/2019 and UI/BD/151552/2021, respectively).

## Acknowledgments

The authors acknowledge the support of the i3S Scientific Platform, HEMS - Histology and Electron Microscopy Service and ALM - Advanced Light Microscopy, both members of the national infrastructure PPBI - Portuguese Platform of Bioimaging (PPBI-POCI-01-0145- FEDER-022122). We Also acknowledge the support of the i3S Proteomics platform, member of the Portuguese Mass Spectrometry Network, integrated in the National Roadmap of Research Infrastructures of Strategic Relevance (ROTEIRO/0028/2013; LISBOA-01-0145-FEDER-022125). The authors also acknowledge the support of the Genomics, Animal facility, and Translational Cytometry i3S Scientific Platform.

All authors approved the final version of the manuscript. This work has not been accepted or published elsewhere.

The authors declare no conflict of interest related to this work.

