## Supplementary Figure for "Increasing the vitamin C transporter SVCT2 in microglia improves synaptic plasticity and restrains memory impairments in Alzheimer’s disease models"

Supplementary Figure 1

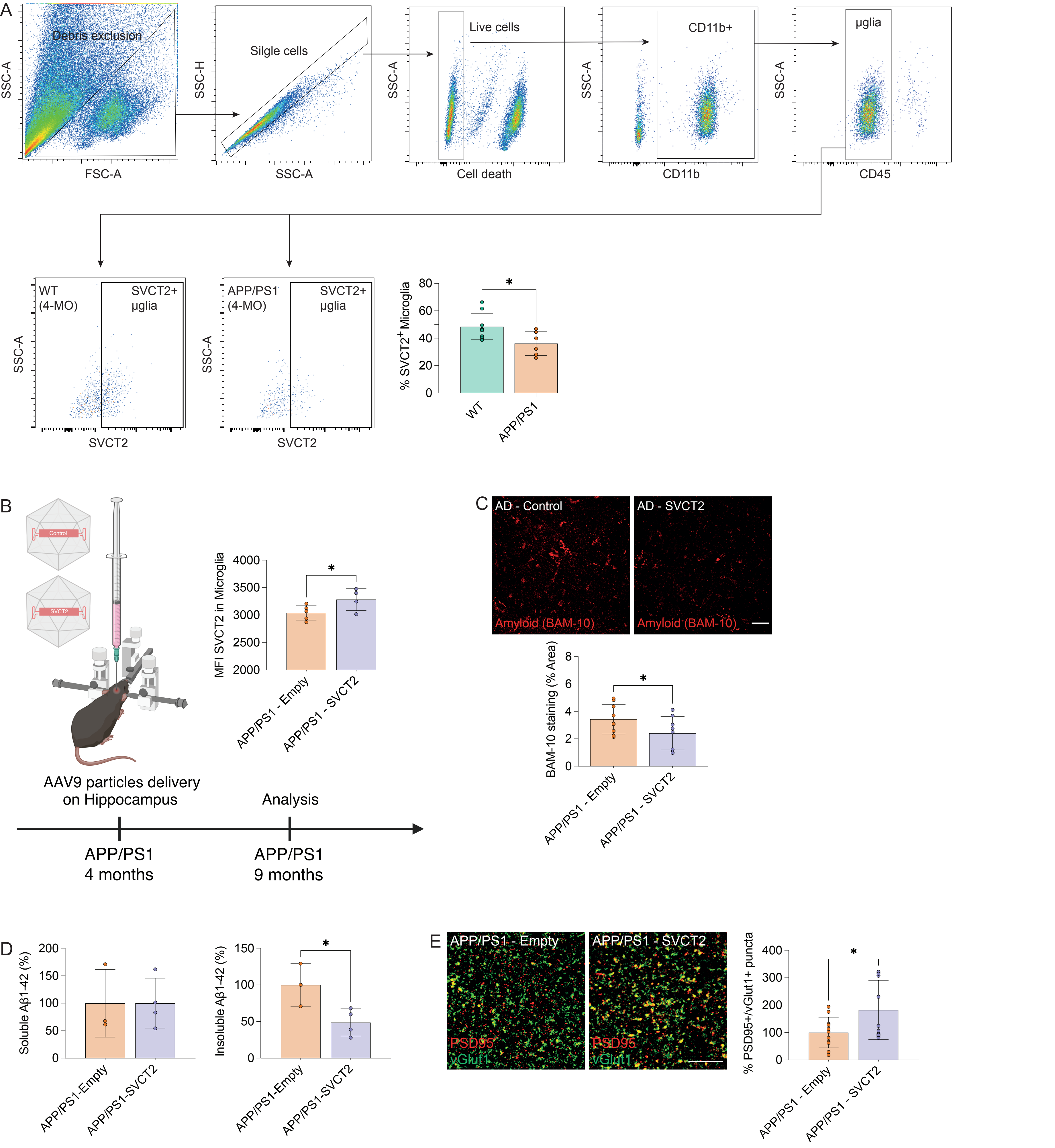

Supplementary Figure 1. SVCT2 overexpression in hippocampal microglia from APP/PS1 mice.

**A: SVCT2 expression in WT and APP/PS1 microglia.** Flow cytometry quantification of $SVCT2^+$ microglia in wild-type (WT) and APP/PS1 transgenic mice at 4-months of age. Utilizing the same gating strategy depicted in Figure 1, the histogram shows the frequency of $SVCT2^+$ microglia within the overall microglial population at 4 months of age. Data are displayed by mean ± SD. *p<0.05 (Unpaired t Test). Each point in the histograms represents a single mouse. WT mice with 4 months of age (N=8); APP/PS1 mice with 4 months of age (N=6).

**B: Schematic depiction of the AAV administration time course in APP/PS1 mice and SVCT2 overexpression.** The strategy was exactly the same as previous described in figure 2 with the exception of the experimental time course. APP/PS1 mice were injected at 4-months of age and analyzed 5 months later (at nine months of age). Flow cytometry was also employed to assess SVCT2 overexpression. The gating strategy was the same described in (A), with the exception that here the result is displayed as the mean fluorescence intensity (MFI), displayed by mean ± SD from both APP/PS1-Empty (N=5) and APP/PS1-SVCT2 (N=4). *p<0.05 (Unpaired t Test). Each point in the histograms represents a single mouse.

**C: Assessment of Amyloid expression in the hippocampus.** Representative confocal images displaying hippocampal BAM-10 staining in APP/PS1 mice transduced with Empty or SVCT2 constructs. Scale bar = 10 µm. Quantitative analysis exhibits the percentage of the BAM-10 stained area plotted as mean ± SD from both APP/PS1-Empty (N=9) and APP/PS1-SVCT2 (N=8). *p<0.05 (Unpaired t Test).

**D: ELISA quantification of $A\beta_{1-42}$ soluble and insoluble forms in the hippocampus.** Soluble and insoluble $A\beta_{1-42}$ levels in detergent (Soluble fraction) and formic acid (insoluble) fractions were analyzed. Graph represents mean ± SD from both APP/PS1-Empty (N=3) and APP/PS1-SVCT2 (N=4). *p<0.05 (Unpaired t Test).

**E: Assessment of excitatory synaptic density in the hippocampus.** Representative confocal images showing postsynaptic density protein 95 (PSD95, red) and presynaptic marker vGlut1 (green) in APP/PS1 mice injected with the control vector (APP/PS1- Empty) compared to APP/PS1 mice overexpressing SVCT2 (APP/PS1 - SVCT2). Colocalization (yellow puncta) depicts excitatory synapses. Graph represents mean ± SD from both APP/PS1-Empty (N=11) and APP/PS1-SVCT2 (N=14). *p<0.05 (Unpaired t Test).

Supplementary table:

Key Resource Table:

| Reagent | Source | Identifier |
|---|---|---|
| **Antibodies** | | |
| Rabbit anti-Ionized Calcium-Binding Adaptor Molecule 1 (Iba-1) | Wako | 016-20001 |
| Chicken anti-mCherry | HenBiotech | HBT008-200 |
| Mouse anti-Beta-Amyloid Protein (Clone: BAM-10) | Merck | A5213 |
| Mouse anti-PSD-95 (Clone:6G6-1C9) | Thermo Fisher Scientific | MA1-045 |
| Rabbit anti vGlut-1 | Synaptic Systems | 135 303 |
| Alexa Fluor™ 647 donkey anti-mouse IgG (H+L) | Thermo Fisher Scientific | A31571 |
| Alexa Fluor™ 488 goat anti-rabbit IgG (H+L) | Thermo Fisher Scientific | A11008 |
| Alexa Fluor™ 568 goat anti-chicken IgY (H+L) | Thermo Fisher Scientific | A11041 |
| Alexa Fluor™ 568 goat anti-mouse IgG (H+L) | Thermo Fisher Scientific | A11004 |
| Alexa Fluor™ 647 goat anti-rabbit IgG (H+L) | Thermo Fisher Scientific | A21244 |
| PE anti-mouse CD45 (Clone 30-F11) | BioLegend | 103106 |
| PE/Cy7 anti-mouse/human CD11b (Clone: M1/70) | BioLegend | 101216 |
| Rabbit anti-SLC23A2 | Abcam | ab229802 |
| Alexa Fluor™ 647 goat anti-rabbit IgG (H+L) | Thermo Fisher Scientific | A21244 |
| Anti-mouse CD16/CD32 Fc blocker | Thermo Fisher Scientific | 14-0161-82 |
| Goat anti SVCT2 | Santa Cruz | G19 |
| **Chemicals, Reagents and Commercial kits** | | |
| Zombie Green™ Fixable kit | BioLegend | 423111 |
| 4',6-diamidino-2-phenylindole, dilactate (DAPI) | Thermo Fisher Scientific | D3571 |
| CD11b MicroBeads human and mouse | Miltenyi Biotec | 130-049-601 |

| | | |
|---|---|---|
| LS columns | Miltenyi Biotec | 130-042-401 |
| RNeasy® Plus Micro Kit | Qiagen | 74034 |
| RNasin® Plus Ribonuclease Inhibitor | Promega | N2611 |
| EASY-Spray™ HPLC Columns | Thermo Fisher Scientific | ES906 |
| Acclaim™ PepMap™ 100 C18 HPLC Columns | Thermo Fisher Scientific | 164946 |
| Hanks' Balanced Salt Solution (HBSS) | Thermo Fisher Scientific | 14170-088 |
| Percoll™ | Merck | 17089102 |
| eBioscience™ Permeabilization Buffer | Thermo Fisher Scientific | 00-8333-56 |
| Pierce™ BCA Protein Assay Kit | Thermo Fisher Scientific | 23225 |
| SuperScript™ VILO™ cDNA Synthesis Kit | Thermo Fisher Scientific | 11754050 |
| Syn-PER™ Synaptic Protein Extraction Reagent | Thermo Fisher Scientific | 87793 |
| Protease inhibitor cocktail | Thermo Fisher Scientific | P8340 |
| Human AB42 Elisa Kit | Thermo Fisher Scientific | KHB3441 |
